# Mechanisms of lactic acid gustatory attraction in *Drosophila*

**DOI:** 10.1101/2021.01.22.427705

**Authors:** Molly Stanley, Britya Ghosh, Zachary F. Weiss, Jaime Christiaanse, Michael D. Gordon

## Abstract

Sour has been studied almost exclusively as an aversive taste modality. Yet, recent work in *Drosophila* demonstrates that specific carboxylic acids are attractive at ecologically relevant concentrations. Here, we demonstrate that lactic acid is an appetitive and energetic tastant, which stimulates feeding through activation of sweet gustatory receptor neurons (GRNs). This activation displays distinct, mechanistically separable, stimulus onset and removal phases. Ionotropic receptor 25a (IR25a) primarily mediates the onset response, which shows specificity for the lactate anion and drives feeding initiation. Conversely, sweet gustatory receptors (Gr64a-f) mediate a non-specific removal response to low pH that primarily impacts ingestion. While mutations in either receptor family have marginal impacts on feeding, lactic acid attraction is completely abolished in combined mutants. Thus, specific components of lactic acid are detected through two classes of receptors to activate a single set of sensory neurons in physiologically distinct ways, ultimately leading to robust behavioural attraction.

## INTRODUCTION

Tastants are canonically classified as belonging to a single taste modality, which is generally sensed by one receptor or family of receptors. However, gustatory detection of some chemical species can be complex, with specific molecular properties differentially acting on multiple receptors. For example, artificial sweeteners can activate both sweet and bitter receptors, NaCl is detected as Na^+^ and Cl^-^ through multiple receptors in different types of gustatory cells, and many bitter compounds inhibit insect sweet receptors (Jaeger et al. 2018; Roebber, Roper, and Chaudhari 2019; Chandrashekar et al. 2010; Behrens, Blank, and Meyerhof 2017; French et al. 2015; Meunier et al. 2003; Jeong et al. 2013; Freeman, Wisotsky, and Dahanukar 2014). Although acids are a particularly diverse class of ligands containing a large variety of side chains in addition to being protonated, how the specific chemical properties of an individual acid influence gustatory detection remains unclear.

Any solution with sufficiently low pH will stimulate aversive gustatory, olfactory, and somatosensory pathways, and acid sensing by the gustatory system, or ‘sour taste’, is traditionally thought to prevent animals from ingesting potentially harmful spoiled or unripe foods (J. Zhang et al. 2019; Charlu et al. 2013; Y. Y. Wang et al. 2011; Chang, Waters, and Liman 2010; Ai et al. 2010; Depetris-Chauvin et al. 2017). Thus, sour has been studied almost exclusively as an aversive taste modality in both mammals and invertebrates (Charlu et al. 2013; Rimal et al. 2019; J. Zhang et al. 2019; Huang et al. 2006). Recently, the proton channel Otop1 was shown to be necessary for low pH detection by mammalian type III taste receptor cells (Teng et al. 2019; J. Zhang et al. 2019; Tu et al. 2018). However, it has been known for a century that ‘sourness’ varies among acids, even at the same pH, suggesting that acid taste involves more than simply the detection of protons (Da Conceicao Neta, Johanningsmeier, and McFeeters 2007; Harvey 1920). This may be particularly relevant for weak acids that are regularly consumed in nutritious foods, including those having undergone fermentation or preservation. Indeed, the addition of acid to foods is known to differentially modify flavor based on the specific acid used (Pfeiffer et al. 2006; Deshpande et al. 2015).

Recent studies in the fruit fly, *Drosophila melanogaster*, demonstrate that individual carboxylic acids differ dramatically in their behavioural valance. While all acids are aversive at high concentrations and low pH, some are attractive at lower concentrations (Rimal et al. 2019). For example, acetic acid and lactic acid can both strongly encourage feeding, likely signaling the presence of energy or beneficial microbes (Qiao et al. 2019). Appetitive responses to acetic acid are starvation-dependent and require sugar-sensing gustatory receptor neurons (GRNs), but the specific receptors involved are unclear (Devineni et al. 2019). Two broadly-expressed ionotropic receptors (IRs), IR25a and IR76b, mediate acid detection during egg laying (Y. Chen and Amrein 2017), but IR76b is not required for appetitive responses to acetic acid, and *IR25a* mutants show only a slight reduction in acetic acid taste (Devineni et al. 2019). Gustatory mechanisms of acid aversion in *Drosophila* have also been elusive until recently, when IR7a was found to mediate rejection of concentrated acetic acid (Rimal et al. 2019). Remarkably, this IR was not involved in attraction to low concentrations of acetic acid, or in the avoidance of other acids. These studies in flies highlight the complexity of acid detection and reaffirm that the specific anion, concentration, and pH can all play a role in sour taste and subsequent feeding behaviour.

To probe the molecular mechanisms of acid detection, we focused on lactic acid attraction. We began with lactic acid as a ligand since it is particularly appetitive to flies (Rimal et al. 2019) and humans report lactic acid as a relatively mild ‘sour’ stimulus (Pfeiffer et al. 2006; Da Conceicao Neta, Johanningsmeier, and McFeeters 2007). Supplementation of *Drosophila* food with lactic acid was previously found to increase lifespan, suggesting that lactic acid consumption is beneficial to flies at concentrations up to 250 mM (Massie and Williams 1979). Moreover, while very little is known about lactic acid taste in insects, lactic acid has been studied extensively as an attractive odorant for mosquito host seeking (McBride 2016; Raji et al. 2019). Interestingly, the anion lactate is also an attractive odorant for rodents, and humans report the smell as ‘sweet’ more than ‘sour’ (Mosienko et al. 2017).

Here, we show that lactic acid acts through sweet taste neurons to strongly stimulate feeding in *Drosophila*. Although lactic acid is also a robust olfactory attractant to flies, olfaction is dispensable for lactic acid feeding. We found that lactic acid produces unique response dynamics in sweet GRNs, which show calcium peaks during both stimulus onset and removal.

Interestingly, the two peaks are mediated by distinct receptor families, with the onset requiring IR25a and the removal response requiring members of the sweet Gustatory Receptor (GR) family. Mutation of either family leaves lactic acid attraction largely intact, suggesting that both onset and removal peaks are salient during feeding. However, flies carrying mutations in both receptor types completely lack attractive lactic acid taste. To our knowledge, this is the first reported example of two co-expressed receptor families mediating distinct physiological responses to a pure chemical in a single sensory neuron type.

## RESULTS

### Lactic acid is an appetitive taste to *Drosophila*

We began by testing lactic acid responses across several different taste-dependent behavioural assays. Consistent with a previous report (Rimal et al. 2019), flies strongly preferred lactic acid over water in a dye-based binary feeding assay, with peak attraction at 250 mM and attraction at all concentrations tested up to 1 M (Fig. 1A). This remained true for mated and virgin *w*^*1118*^ females, *w*^*1118*^ males, and *Canton S* females and males (Fig. S1A). To ensure that lactic acid attraction was not due to any effects of acid on texture of the agar-based food in this assay, we also performed a Capillary Feeder (CAFE) assay, which uses liquid solutions, and found a strong preference for 250 mM lactic acid maintained over 24 hours (Fig. S1B). We also quantified the proboscis extension reflex (PER), which measures the acute appetitiveness of taste stimuli.

**Figure 1:**
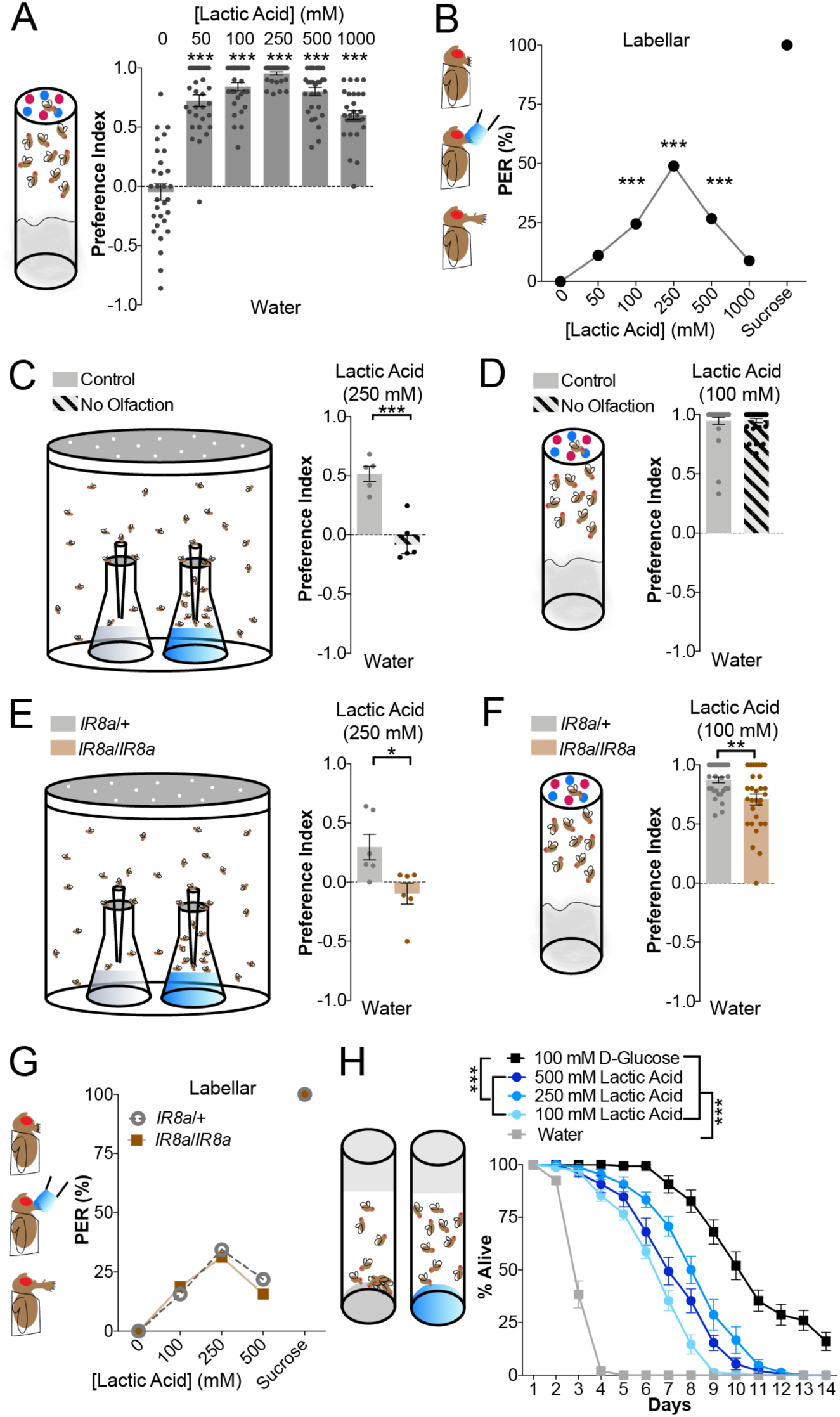
Lactic acid is an appetitive taste to *Drosophila*. (**A**) Schematic of the dye-based binary feeding assay (left) and lactic acid attraction in control *w*^*1118*^ flies using this assay (right). Positive values indicate preference for lactic acid at the indicated concentration; negative values indicate preference for water. Bars represent mean ±SEM. n=30 groups of 10 flies each per concentration. Asterisks denote significant difference from negative control (0 mM, water) by one-way ANOVA with Dunnett’s post test, ∗∗∗p<.001. (**B**) Schematic of labellar PER assay (left) and lactic acid PER in control *w*^*1118*^ flies using this assay (right). n=45 flies and dots represent the mean. 500 mM sucrose was used as a positive control. Asterisks denote significant difference from negative control (0 mM, water) by one-way ANOVA with Dunnett’s post test, ∗∗∗p<.001. (**C**) Schematic of the trap olfactory assay (left) and lactic acid attraction in control *w*^*1118*^ flies with and without olfaction (removal of antennae and maxillary palps) using this assay (right). Positive values indicate preference for 250 mM lactic acid; negative values indicate preference for water. Bars represent mean ±SEM. n=5 sets of 40 flies each per group. Asterisks denote significance between groups by unpaired t-test, ∗∗∗p<.001. (**D**) Preferences of control *w*^*1118*^ flies with and without olfaction (removal of antennae and maxillary palps) in the dye-based binary feeding assay. Positive values indicate preference for 100 mM lactic acid; negative values indicate preference for water. Bars represent mean ±SEM. n=30 groups of 10 flies each per group, no significant differences by unpaired t-test. (**E**) Preferences of *IR8a* mutants and heterozygous controls in the trap olfactory assay. Positive values indicate preference for 250 mM lactic acid; negative values indicate preference for water. Bars represent mean ±SEM. n=6 sets of 40 flies each per genotype. Asterisks denote significance between groups by unpaired t-test, ∗p<.05. (**F**) Preferences of *IR8a* mutants and heterozygous controls in the binary feeding assay. Positive values indicate preference for 100 mM lactic acid; negative values indicate preference for water. Bars represent mean ±SEM. n=30 groups of 10 flies per genotype. Asterisks denote significance between groups by unpaired t-test, ∗∗p<.01. (**G**) Labellar PER of *IR8a* mutants and heterozygous controls. n=32 flies per genotype and dots represent the mean. 500 mM sucrose was used as a positive control. No significant differences by two-way ANOVA with Tukey’s post test. (**H**) Survival of control *w*^*1118*^ flies on indicated solutions. Points represent mean ±SEM. n=15 groups of 10 flies per solution plotted as % of flies alive per group per day. Asterisks denote significance between groups by two-way ANOVA with Tukey’s post test, ∗∗∗p<.001.

Stimulation of labellar taste sensilla with lactic acid produced dose-dependent PER that mirrored the binary feeding assay, with significant responses at 100-500 mM and a maximum response around 250 mM (Fig. 1B). Tarsal PER to lactic acid was weak but significant from concentrations of 100 mM to 500 mM (Fig. S1C).

Considering that lactic acid is an attractive olfactory cue to other insects, we wanted to clarify the role of olfaction versus gustation in lactic acid feeding. In an olfactory trap assay, control flies showed a clear preference for 250 mM lactic acid, but this preference was completely abolished following surgical removal of the olfactory organs (Fig. 1C). However, the same surgery had no effect on lactic acid preference and consumption in the binary feeding assay, indicating that olfaction is dispensable for lactic acid feeding attraction (Fig. 1D). As independent verification, we also tested flies with mutated *IR8a*, which mediates lactic acid olfactory attraction in *Aedes aegypti* (Raji et al. 2019) and olfactory acid aversion in *Drosophila* (Ai et al. 2013). *IR8a* mutants showed no olfactory attraction to lactic acid, but maintained strong preference in the binary feeding assay (Fig. 1E,F). Although this preference was slightly reduced in *IR8a* mutants compared to controls, labellar (Fig. 1G) and tarsal (Fig. S1D) PER responses were normal. Therefore, we conclude that appetitive taste and feeding responses to lactic acid are independent of olfaction.

To investigate a potential reason for flies’ strong attraction to lactic acid, we quantified the ability of lactic acid to provide energy. We found that lactic acid presented as the sole energy source significantly promoted survival, although to a lesser extent than D-glucose (Fig. 1H). Thus, lactic acid is an appetitive, attractive, and energetic compound for *Drosophila*.

### Sweet GRNs are necessary for lactic acid feeding attraction

To examine the cellular basis of attractive lactic acid taste, we used Kir2.1 expression to systematically silence five distinct GRN classes that encompass almost every taste neuron on the fly labellum (Jaeger et al. 2018). Only the silencing of sweet GRNs, labeled by *Gr64f-Gal4*, abolished lactic acid attraction (Fig. 2A). We found that Gr64f GRNs are absolutely required for appetitive responses to lactic acid in both binary choice feeding and PER (Fig. 2B,C). Notably, flies lacking sweet taste showed concentration-dependent avoidance of lactic acid, suggesting that lactic acid stimulates a parallel aversive pathway (Fig. 2B). While *Gr64f-Gal4* is the standard for driving expression in all sweet GRNs, it is also expressed in Olfactory Receptor Neurons (ORNs) projecting to the antennal lobe (Menuz et al. 2014)(Fig. S2A). To confirm that the feeding effects were due to the gustatory expression of *Gr64f-Gal4* in the subesophageal zone (SEZ) and not an effect on olfaction, we repeated the olfactory trap assay with Gr64f-silenced flies and found no change in lactic acid attraction (Fig. S2B). We also silenced a majority of sweet GRNs using *Gr64e-Gal4*, which has no olfactory expression (Fig. S2C), and saw elimination of feeding attraction to lactic acid (Fig. S2D).

**Figure 2:**
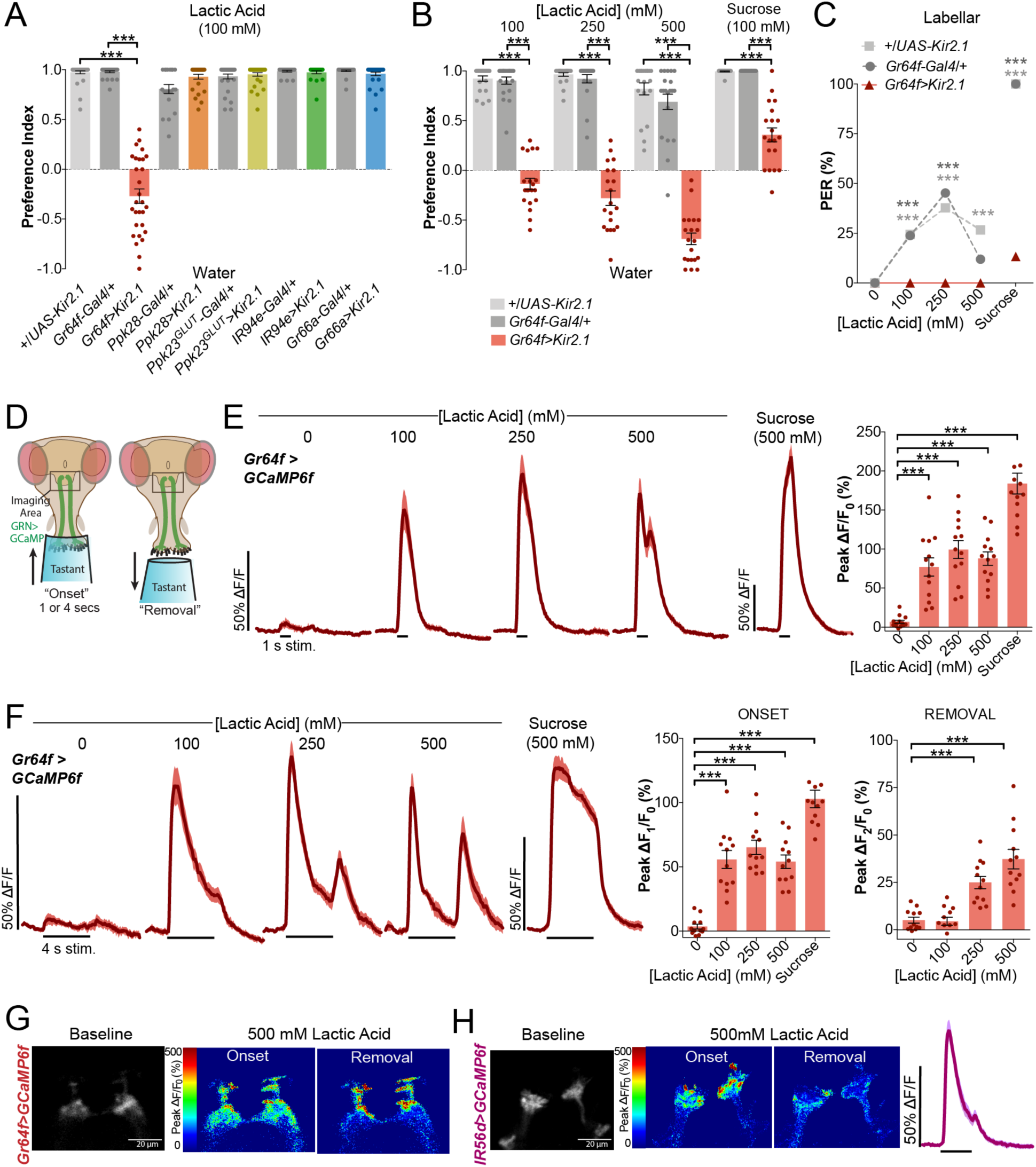
Sweet GRNs are necessary for lactic acid feeding attraction. (**A**) Preferences for lactic acid in the binary feeding assay, following silencing of distinct classes of GRNs using Kir2.1 (Gr64f= ‘sweet’, Ppk28= ‘water’, Ppk23^GLUT^= ‘high salt’, IR94e= orphan GRN, Gr66a= ‘bitter’). Positive values indicate preference for 100 mM lactic acid; negative values indicate preference for water. Bars represent mean ±SEM. n=20-36 groups of 10 flies per genotype. Asterisks denote significant differences by one-way ANOVA with Tukey’s post test, ∗∗∗p<.001. (**B**) Preferences in the binary feeding assay following sweet GRN silencing. Positive values indicate preference for lactic acid or sucrose at indicated concentrations; negative values indicate preference for water. Bars represent mean ±SEM. n=20 groups of 10 flies per genotype per concentration. Asterisks denote significance by two-way ANOVA with Tukey’s post test, ∗∗∗p<.001. (**C**) Labellar PER of flies with sweet GRNs silenced and genotype controls. n=42-45 flies per genotype and dots represent the mean. 500 mM sucrose was used as a positive control. Asterisks denote significance between genotypes by two-way ANOVA with Tukey’s post test, ∗∗∗p<.001. (**D**) Schematic of *in vivo* calcium imaging preparation. Labellar taste neurons are stimulated with tastant while GCaMP6f fluorescence is recorded in the synaptic terminals in the SEZ region of the brain. Tastant is left over the labellum for either 1 or 4 s. (**E**) Sweet GRN calcium responses of *Gr64f>GCaMP6f* flies to 1-second stimulations. Lines and shaded areas represent mean ±SEM over time (left). Peak fluorescence changes during each stimulation (right). Bars represent mean ±SEM. n=13 flies. Asterisks denote significant difference from negative control (0 mM, water) by one-way ANOVA with Dunnett’s post test, ∗∗∗p<.001. (**F**) Sweet GRN calcium responses to 4-second stimulations. Lines and shaded areas represent mean ±SEM over time (left). Quantification of ‘onset’ and ‘removal’ peak fluorescence changes during each stimulation and removal of stimulus (right). Bars represent mean ±SEM. n=12 flies. Asterisks denote significant difference from negative control (0 mM, water) by one-way ANOVA with Dunnett’s post test, ∗∗∗p<.001. (**G**) Representative heat map showing *Gr64f>GCaMP6f* fluorescence changes with 500 mM lactic acid stimulus onset vs. removal. (**H**) Representative heat map showing *IR56d>GCaMP6f* fluorescence changes with 500 mM lactic acid stimulus onset vs. removal. Right: GCaMP6f fluorescence changes over time with 500 mM lactic acid stimulation, lines and shaded area represent mean ±SEM. n=15 flies.

Next, we measured sweet GRN calcium responses to lactic acid by recording GCaMP6f fluorescence in Gr64f axon terminals of the SEZ (Fig. 2D). As expected, standard 1-second stimulations (Jaeger et al. 2018) revealed strong calcium responses to lactic acid (Fig. 2E). However, we noted an emergence of two peaks at 500 mM: one coinciding with the onset of stimulation and one with stimulus removal. To investigate these unusual calcium kinetics further, we repeated the imaging experiment with 4-second stimulations to allow better separation of the two peaks. This revealed a distinct removal peak that appeared at concentrations of 250 mM and above (Fig. 2F). Unlike lactic acid, sucrose stimulation did not produce two peaks, in agreement with a recent report investigating calcium kinetics in GRNs (Devineni et al. 2020). Interestingly, the two lactic acid peaks appeared somewhat spatially distinct in the SEZ, with removal activity primarily localized in dorsal Gr64f projections (Fig. 2G). The onset peak region appeared to overlap with the subset of fatty-acid sensitive sweet neurons labelled by *IR56d-Gal4* (Tauber et al. 2017) (IR56d+), whereas the removal peak region appeared to overlap with an IR56d negative subset (IR56d-). Indeed, the IR56d subset of sweet GRNs exhibit robust onset responses to 500 mM lactic acid, with little to no removal peak (Fig. 2H).

Overall, these results show that lactic acid strongly activates sweet GRNs with unique calcium kinetics, corresponding to both stimulus onset and removal, and this sweet GRN activity is necessary for the appetitive responses and feeding preference for lactic acid.

### pH influences both onset and removal calcium peaks in GRNs

The pH of 100 mM lactic acid is normally about 3, while 500 mM concentrations have a pH of about 2. Since emergence of the sweet GRN removal calcium peak is correlated with the lower pH of higher acid concentrations, we next investigated the contribution of pH to these calcium kinetics. We lowered the pH of 100 mM lactic acid to 2 by addition of HCl, and raised it to 7 using NaOH. To control for the addition of HCl, we included a stimulation of HCl alone at an equivalent pH of 2, and to control for the Na^+^ in NaOH, we added an equivalent concentration of NaCl into the 100 mM lactic acid control. In sweet GRNs, we found that HCl produced a minimal onset peak and a prominent removal peak (Fig. 3A). Similarly, pH 2 lactic acid evoked a lower onset peak and stronger removal peak compared with control lactic acid, exhibiting kinetics more closely resembling responses to higher acid concentrations (Fig. 3A). Conversely, 100 mM neutral lactic acid produced a weaker onset peak and no removal peak. Although the addition of Na^+^ during pH adjustment likely contributes to the onset peak, the lack of removal peak is consistent with lactic acid removal responses being pH-dependent. Based on these experiments, we posited a model in which two distinct chemical properties of lactic acid act on sweet GRNs: lactate anions evoke a strong, specific onset response, while acidity contributes to onset responses and also produces prominent removal peaks (Fig. 3B).

**Figure 3:**
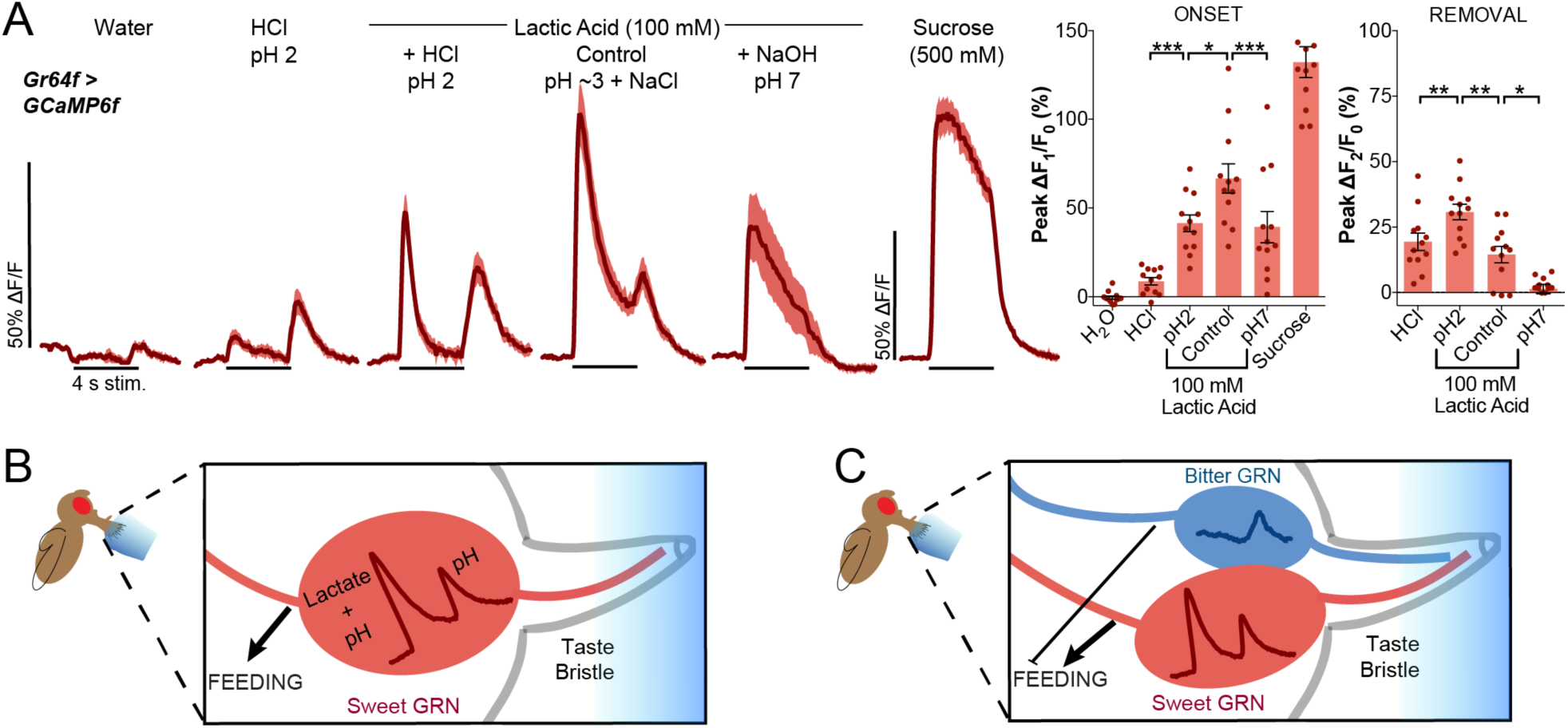
pH influences both onset and removal calcium peaks in GRNs. (**A**) Sweet GRN calcium responses to pH-adjusted 100 mM lactic acid and control solutions. Lines and shaded areas represent mean ±SEM (left). Quantification of ‘onset’ and ‘removal’ peak fluorescence changes during each stimulation and removal of stimulus (right). Bars represent mean ±SEM over time. n=12 flies, each receiving all tastants in random order (except water always first and sucrose last). Asterisks denote significant differences between control 100 mM lactic acid and other test solutions by one-way ANOVA with Sidak’s post test, ∗p<.05, ∗∗p<.01,∗∗∗p<.001. (**B**) Model of a single sweet GRN inside a taste bristle representing how sweet GRNs respond to lactic acid stimulation: the onset calcium peak is largely in response to lactate but low pH also contributes, whereas the removal calcium peak is generated by a change in pH. (**C**) Model of sweet and bitter GRNs within a taste bristle simultaneously responding to lactic acid stimulation and their combined influence on feeding behaviour.

Previous work has demonstrated that the low pH of other carboxylic acids can activate bitter GRNs (Charlu et al. 2013). Therefore, we measured bitter GRN responses to lactic acid for comparison (Fig. S3A). Short (1 s) lactic acid stimulations produced dose-dependent responses with kinetics similar to bitter stimulations (Fig. S3B). However, longer (4 s) stimulations revealed that lactic acid primarily produces calcium peaks upon stimulus removal (Fig. S3C,D). At the highest concentration tested (500 mM), calcium levels gradually increase during the 4 s stimulation, peak with removal, and remain elevated (Fig. S3D). Stimulating with the panel of pH-adjusted solutions, we find that HCl alone produces low, sustained responses beginning with stimulus onset (Fig. S3E). pH 2 lactic acid evokes strong onset responses, with calcium levels remaining high even after stimulus removal. Since NaCl alone can activate bitter GRNs (Jaeger et al. 2018), its addition to the pH 3 lactic acid control undoubtedly contributes to the onset peak not seen from pure 100 mM lactic acid. However, most interestingly, neutral lactic acid produced no response in bitter GRNs, despite the presence of Na^+^ (Fig. S3E).

The co-activation of sweet and bitter GRNs by lactic acid is consistent with our behavioural experiments revealing dose-dependent aversion in the absence of sweet GRN function (Fig. 2B). Thus, stimulation of sweet GRNs overrides bitter GRN activity to drive lactic acid feeding (Fig. 3C). Since we were most interested in the mechanisms of lactic acid attraction, we focused entirely on sweet GRN responses from this point forward.

### IR25a partially mediates lactic acid attraction

We initially took a candidate receptor approach to uncovering the molecular mechanisms of acid detection, beginning with IR25a. IR25a is a broadly expressed co-receptor required for most IR-mediated taste detection, including acid detection by tarsal ‘sour’ GRNs during oviposition (Jaeger et al. 2018; Lee et al. 2018; Ahn, Chen, and Amrein 2017; Sánchez-Alcañiz et al. 2018). Mild defects in acetic acid PER have been reported for *IR25a* mutants (Devineni et al. 2019), and IR25a was a candidate receptor for propionic acid feeding attraction in *Drosophila* larvae (Depetris-Chauvin et al. 2017). We found that *IR25a* mutants display a strong but incomplete reduction in lactic acid PER (Fig. 4A). However, these mutants show only a slight reduction in preference for lactic acid in the binary feeding assay (Fig. 4B). Since IR25a is broadly expressed in chemosensory neurons, we rescued IR25a specifically in sweet GRNs and found that this restored normal lactic acid feeding preference (Fig. 4B). These data indicate that flies possess IR25a-dependent and IR25a-independent mechanisms for gustatory attraction to lactic acid, and that IR25a appears to play a more prominent role in the acute feeding initiation response measured by PER. We also examined IR76b, which is another broad IR co-receptor with many functions that overlap with IR25a; however, *IR76b* mutants had no reduction in lactic acid PER or feeding preferences (Fig. 4C,D). In fact, we observed marginally increased attraction at some concentrations (Fig. 4D).

**Figure 4:**
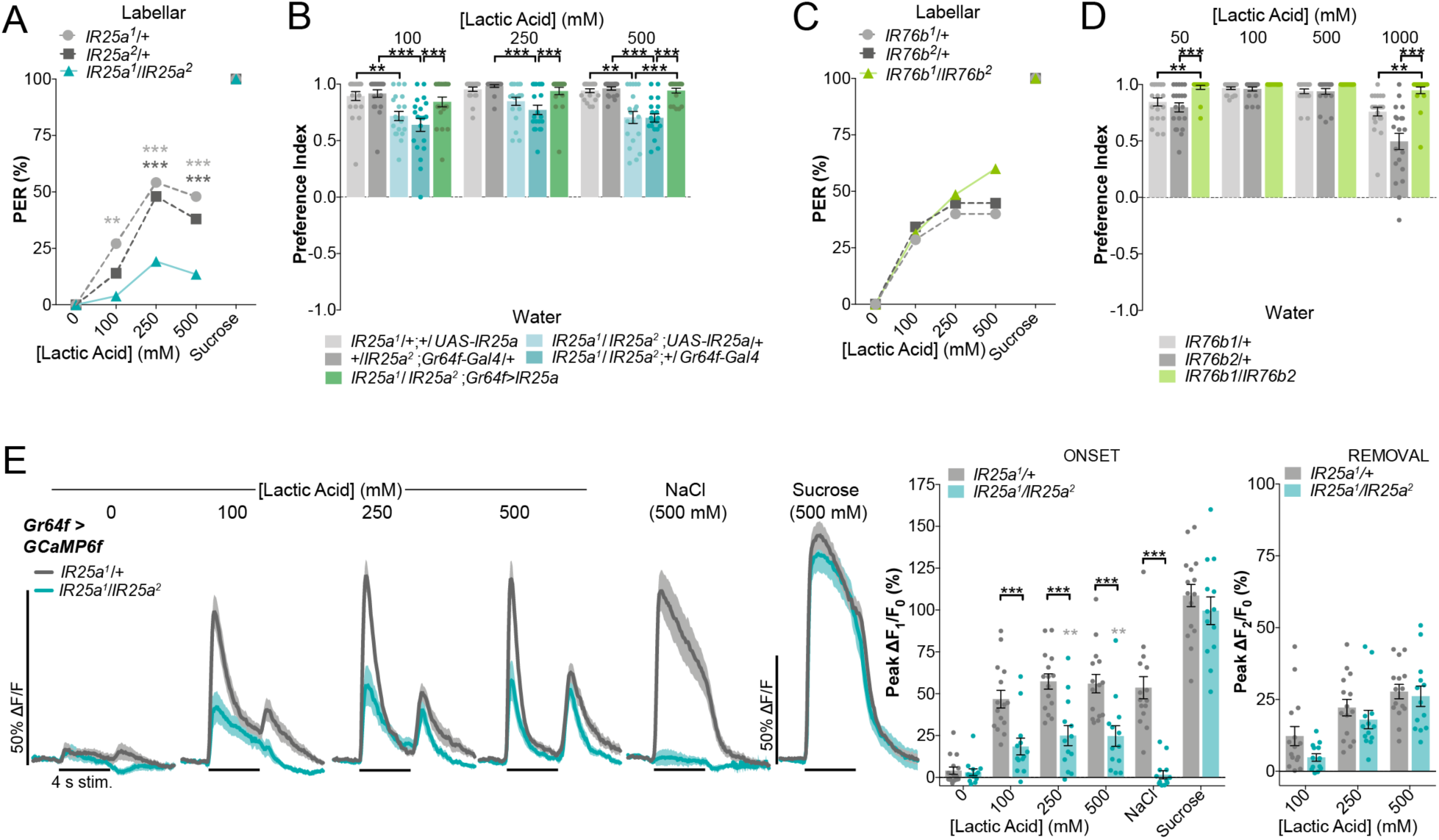
IR25a partially mediates lactic acid attraction. (**A**) Labellar PER of *IR25a* mutants and heterozygous controls. n=48-52 flies per genotype and dots represent means. 500 mM sucrose was used as a positive control. Asterisks denote significant differences by two-way ANOVA with Tukey’s post test, ∗∗p<.01,∗∗∗p<.001. (**B**) Preferences of *IR25a* mutants, flies with rescued IR25a expression in sweet GRNs, and genotype controls in the binary feeding assay. Positive values indicate preference for lactic acid at indicated concentrations; negative values indicate preference for water. Bars represent mean ±SEM. n=20 groups of 10 flies per genotype per concentration. Asterisks denote significance between by two-way ANOVA with Tukey’s post test, ∗∗p<.01,∗∗∗p<.001. (**C**) Labellar PER of *IR76b* mutants and heterozygous controls. n=35-38 flies per genotype and dots represent means. No significant differences by two-way ANOVA with Tukey’s post test. (**D**) Preferences of *IR76b* mutants and heterozygous controls in the binary feeding assay. Positive values indicate preference for lactic acid at indicated concentrations; negative values indicate preference for water. Bars represent mean ±SEM. n=20 groups of 10 flies per genotype per concentration. Asterisks denote significance between by two-way ANOVA with Tukey’s post test, ∗∗p<.01,∗∗∗p<.001. (**E**) Sweet GRN calcium responses in *IR25a* mutants and heterozygous controls with 4-second stimulations. Lines and shaded areas represent mean ±SEM over time (left). Quantification of ‘onset’ and ‘removal’ peak fluorescence changes during each stimulation and removal of stimulus (right). Bars represent mean ±SEM. n=13-15 flies per genotype. Black asterisks denote significant differences by two-way ANOVA with Sidak’s post test, ∗∗∗p<.001. Grey asterisks denote significance within mutants by additional one-way ANOVA with Dunnett’s post test comparing test solutions to water, ∗∗p<.01.

Consistent with our behavioural data, calcium imaging of *IR25a* mutants revealed partial, but significant, reductions in sweet GRN responses to 1 s lactic acid stimuli, which were rescued by cell-type specific expression of *IR25a* (Fig. S4A). To more closely investigate the calcium kinetics, we performed calcium imaging with 4-second stimulations and observed significantly reduced onset peaks in *IR25a* mutants at all concentrations of lactic acid, but no effect on the removal peaks (Fig. 3E). This partial effect contrasts with the total loss of NaCl responses and normal sucrose-evoked activity observed in the same *IR25a* mutants (Jaeger et al. 2018) (Fig. 4E, S4A). Qualitatively, the reduction in onset responses appeared primarily in the IR56d+ projection area, leaving residual onset and normal removal responses in the IR56d-region of the SEZ (Fig. S4B). Moreover, although salt activity in sweet GRNs is *IR25a*-dependent, stimulations with NaCl do not produce a removal peak, demonstrating that two peaks is not a general property of IR25a activation (Fig. 4E). Together, our results suggest a model in which IR25a mediates sweet GRN onset responses to the lactate anion, while an independent mechanism drives non-specific onset and removal responses to acidic stimuli.

### Sweet Gustatory Receptors partially mediate lactic acid attraction

We next sought to uncover the mechanism underlying IR25a-independent lactic acid responses in sweet GRNs. A screen of other IRs, including IR56d, which is responsible for fatty acid taste (Sánchez-Alcañiz et al. 2018), revealed no defects in lactic acid preference (Fig. S5A).

Therefore, we turned to gustatory receptors (GRs), which represent the other major class of receptors found in sweet GRNs and are known to detect various sugars (Yavuz et al. 2014; Jiao et al. 2008; Dahanukar et al. 2007; Miyamoto et al. 2012). Surprisingly, every sugar GR mutant we tested showed a significant reduction in lactic acid attraction (Fig. 5A). However, the strongest phenotype appeared to be in *Gr64a*^*2*^ mutants, which have a deletion covering *Gr64a, Gr64b*, and *Gr64c*. Testing an independent deletion of the entire Gr64 cluster of six sweet GRs (Δ*Gr64a-f*) produced an equivalent phenotype, which was not enhanced by further removal of the three remaining sweet GRs (Δ*8* sugar GRs, *Gr43a*^*LEXA*^). In order to probe the role of GRs in lactic acid taste we continued by primarily studying the Δ*Gr64a-f* deletion because it combined a strong phenotype with access to genetic manipulations that are unfeasible in flies lacking all nine sweet GRs.

**Figure 5:**
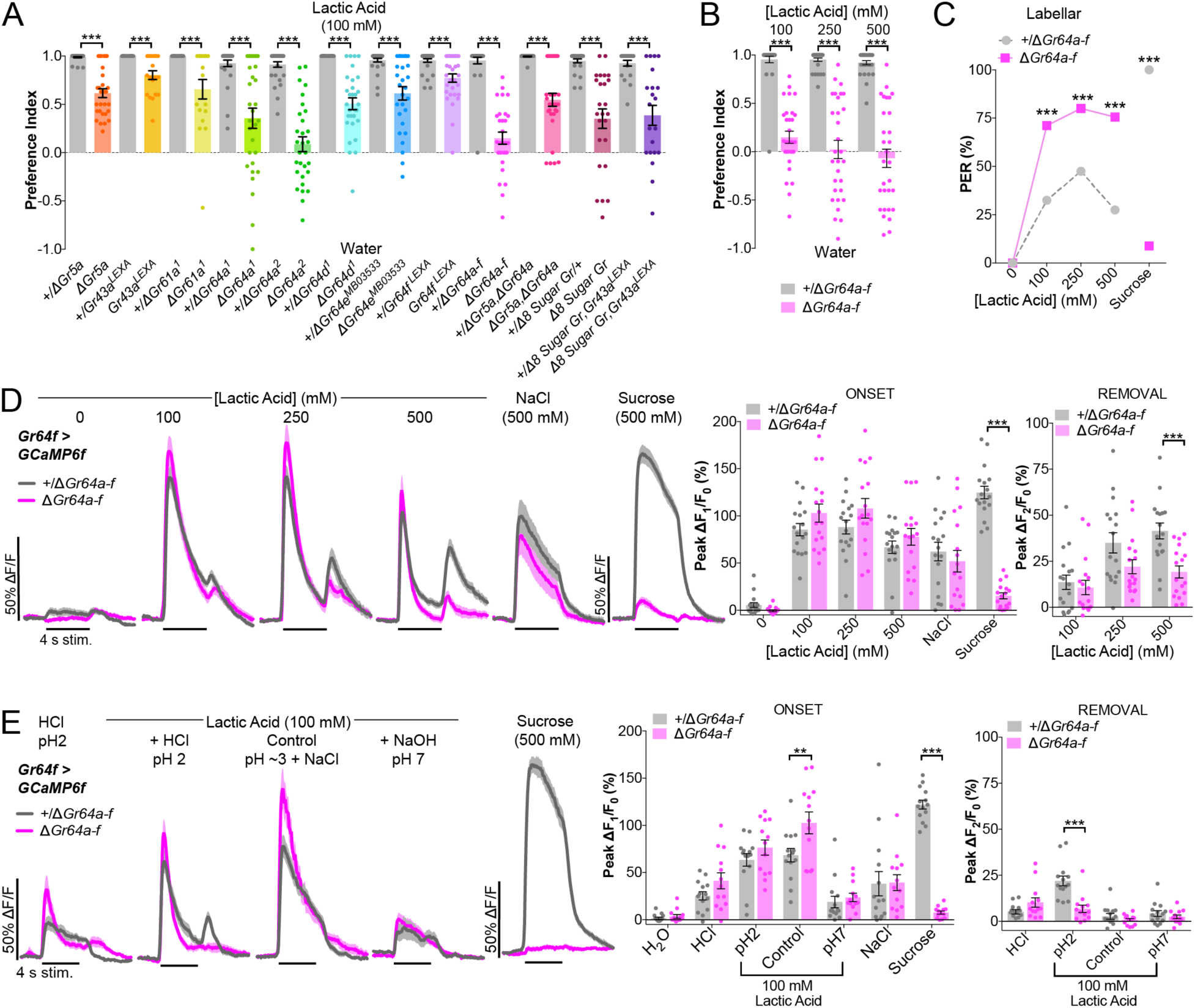
Sweet Gustatory Receptors partially mediate lactic acid attraction. (**A**) Binary choice feeding preferences of flies with single or combined sweet GR mutations (colors) or heterozygous controls (grey). Positive values indicate preference for 100 mM lactic acid; negative values indicate preference for water. Bars represent mean ±SEM. n=18-30 groups of 10 flies per genotype (group sizes matched for each mutant and control). Asterisks indicate significant differences by unpaired t-test for each mutant vs. heterozygous control, ∗∗∗p<.001. (**B**) Binary choice feeding preferences of Δ*Gr64a-f* mutants or heterozygous controls. Positive values indicate preference for lactic acid at indicated concentrations; negative values indicate preference for water. Bars represent mean ±SEM. n=30 groups of 10 flies per genotype per concentration. Asterisks denote significance between by two-way ANOVA with Tukey’s post test, ∗∗∗p<.001. (**C**) Labellar PER of Δ*Gr64a-f* mutants or heterozygous controls. n=40-45 flies per genotype and dots represent the mean. 500 mM sucrose was used as a positive control in heterozygotes. Asterisks denote significance between by two-way ANOVA with Tukey’s post test, ∗∗∗p<.001. (**D**) Sweet GRN calcium responses in Δ*Gr64a-f* mutants and heterozygous controls with 4-second stimulations. Lines and shaded areas represent mean ±SEM over time (left). Quantification of ‘onset’ and ‘removal’ peak fluorescence changes during each stimulation and removal of stimulus (right). Bars represent mean ±SEM. n=17 flies per genotype. Asterisks denote significant differences by two-way ANOVA with Sidak’s post test, ∗∗∗p<.001. (**E**) Sweet GRN calcium responses to with pH-adjusted 100 mM lactic acid and control solutions. Lines and shaded areas represent mean ±SEM over time (left). Negative and positive control curves (water, NaCl) are not shown due to space restrictions, the kinetics were not qualitatively different from (E). Quantification of ‘onset’ and ‘removal’ peak fluorescence changes during each stimulation and removal of stimulus (right). Bars represent mean ±SEM. n=13-14 flies per genotype. Asterisks denote significant differences between genotypes by two-way ANOVA with Sidak’s post test, ∗∗p<.01,∗∗∗p<.001.

Δ*Gr64a-f* mutants have a strongly reduced feeding preference for lactic acid across concentrations from 100 mM to 500 mM (Fig. 5B). However, these mutants do not show the clear switch to behavioural aversion evident with sweet GRN silencing (Fig. 2B), highlighting the existence of a GR-independent pathway for lactic acid attraction. Strikingly, we found that Δ*Gr64a-f* mutants have substantially elevated PER to lactic acid (Fig. 5C), consistent with a prior report showing enhanced PER to acetic acid in sweet GR mutants (Devineni et al. 2019). This presents a paradoxical mismatch between the apparent role of sweet GRs in the binary feeding assay and PER.

In an attempt to reconcile these opposing behavioural results, we performed calcium imaging of sweet GRNs in Δ*Gr64a-f* mutants. As expected, control stimulations showed a significant reduction in sucrose responses with no impact on NaCl (Fig. 5D, Fig. S5C). Although 1-second stimulations with lactic acid revealed no significant difference between Δ*Gr64a-f* mutants and controls (Fig. S5C), 4-second stimulations produced two trends: the onset peaks trended higher in the mutants, and the removal peaks were lower with significance at 500 mM lactic acid (Fig. 5D). Qualitatively, the localization of onset and removal responses in SEZ projections was also less separable in Δ*Gr64a-f* mutants, with the IR56d-region of the SEZ largely inactive in the mutants (Fig. S5B).

Since GR mutations affect the pH-sensitive removal response to lactic acid, we next investigated the interaction of GRs and pH by performing pH-adjusted 100 mM lactic acid stimulations in Δ*Gr64a-f* mutants. Strikingly, Δ*Gr64a-f* mutants completely lacked the removal peak evoked by pH 2 lactic acid (Fig. 5E). Moreover, while we observed a significant increase in the onset response to the lactic acid control in Δ*Gr64a-f* mutants, this enhancement was absent in the response to neutral lactic acid (Fig. 5E). These results suggest a role for sweet GRs that is opposite to that of IR25a: GRs mediate the second peak to acid removal, and have a minor effect on limiting the onset peak. The GR-mediated removal response also appears sensitive to GR dose, as heterozygous controls lacked a removal peak to HCl alone (Fig. 5E).

Taken together, our GR mutant imaging and behavioural data suggest that GR-mediated removal responses are dispensable for feeding initiation (PER), but play an important role in feeding programs that drive consumption.

### Lactic acid attraction is abolished in combined mutants for IR25a and sweet GRs

Having uncovered two receptor types that are each partially responsible for behavioural attraction to lactic acid and mediate distinct calcium responses in sweet GRNs, we next generated flies that had mutations in both *IR25a* and Δ*Gr64a-f*. In these combined *IR25a*, Δ*Gr64a-f* mutants, sweet GRN activation by lactic acid was completely abolished (Fig. 6A,B). While NaCl responses were also lost due to the *IR25a* mutation, sucrose responses were substantially reduced but not completely eliminated (Fig. 6A,B), providing evidence that the GRNs were still intact and able to respond to stimulation. Consistent with the observed physiology, labellar PER to lactic acid was completely eliminated in combined mutants (Fig. 6C), similar to sweet GRN silencing (Fig. 2C). Moreover, lactic acid preference in the binary choice feeding assay was also fully eliminated, leading to behavioural aversion (Fig. 6D) that is equivalent or stronger than with sweet GRN silencing (Fig. 2B). Flies that were homozygous for one receptor mutation and heterozygous for the other receptor mutation showed feeding preferences similar to the individual mutants: Δ*Gr64a-f* mutants had a stronger phenotype than *IR25a* mutants but all were still attracted to lactic acid to some extent (Fig. 5D). These experiments confirm that appetitive lactic acid detection is mediated by two receptor families that each contribute to distinct components of the physiological response, and that these responses are only abolished upon removal of both receptor types.

**Figure 6:**
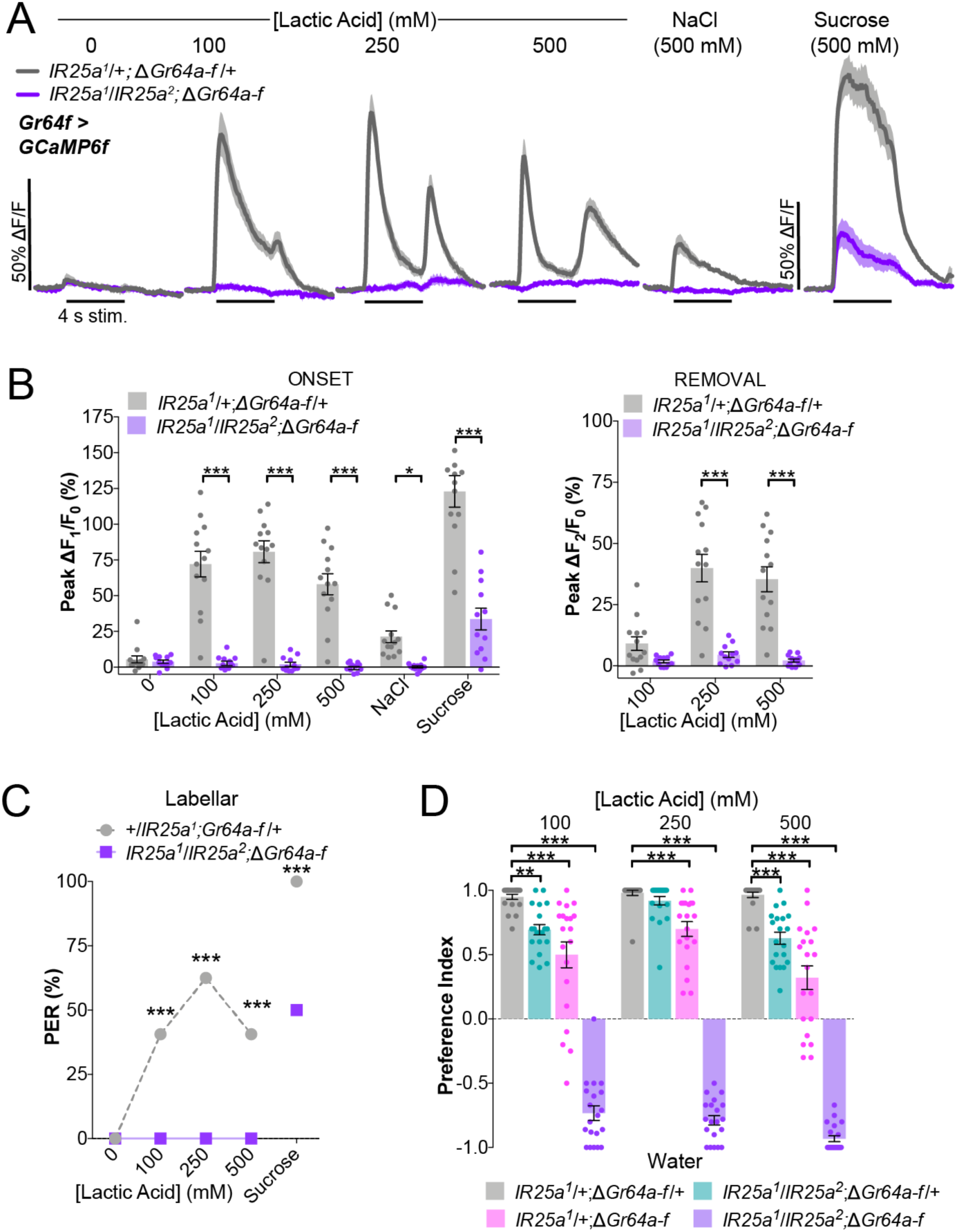
Lactic acid attraction is abolished in combined mutants for IR25a and sweet GRs. (**A**) Sweet GRN calcium responses in *IR25a*, Δ*Gr64a-f* mutants and heterozygous controls with 4-second stimulations. Lines and shaded areas represent mean ±SEM over time. (**B**) Quantification of ‘onset’ and ‘removal’ peak fluorescence changes during each stimulation and removal of stimulus. Bars represent mean ±SEM. n=12-13 flies per genotype. Asterisks denote significant differences by two-way ANOVA with Sidak’s post test, ∗p<.05, ∗∗∗p<.001. (**C**) Labellar PER of *IR25a*, Δ*Gr64a-f* mutants or heterozygous controls. n=32 flies per genotype and dots represent the mean. 500 mM sucrose was used as a positive control in heterozygotes. Asterisks denote significance between by two-way ANOVA with Tukey’s post test, ∗∗∗p<.001. (**D**) Binary choice feeding preferences of *IR25a*, Δ*Gr64a-f* mutants or heterozygous controls. Positive values indicate preference for lactic acid at indicated concentration; negative values indicate preference for water. Bars represent mean ±SEM. n=20 groups of 10 flies per genotype. Asterisks indicate significant differences by two-way ANOVA with Dunnett’s post test, ∗∗p<.01, ∗∗∗p<.001.

### Differentiation between lactic and other attractive acids requires IR25a

Our model for attractive lactic acid taste posits that IR25a primarily detects the lactate anion while sweet GRs more generally respond to low pH. One prediction of this model is that sweet GR mutants should still prefer lactic acid over less attractive carboxylic acids, but *IR25a* mutants should lose this distinction. We chose to test this idea using acetic and propionic acid based on previous reports (Devineni et al. 2019; Rimal et al. 2019; Depetris-Chauvin et al. 2017), and began by confirming that 100 mM concentrations of both acids are attractive compared to water, but less so than lactic acid (Fig. S6A). In line with this feeding preference, acetic and propionic acid elicit calcium responses in sweet GRNs that are relatively weaker than lactic acid (Fig. S6B).

As expected, *IR25a* mutants show only minor reductions in preference for all three acids compared to water in the binary choice assay (Fig. 7A). To accurately measure relative attraction to the three acids, we next tested preference between 200 mM concentrations of lactic acid and each of the other two acids, all with a pH of 2. For this experiment, we employed an acute CAFE (volume-based) binary choice assay because pH-adjusted solutions can be used without any modification, such as the addition of agar or dye, that could interact with the acids differently.

**Figure 7:**
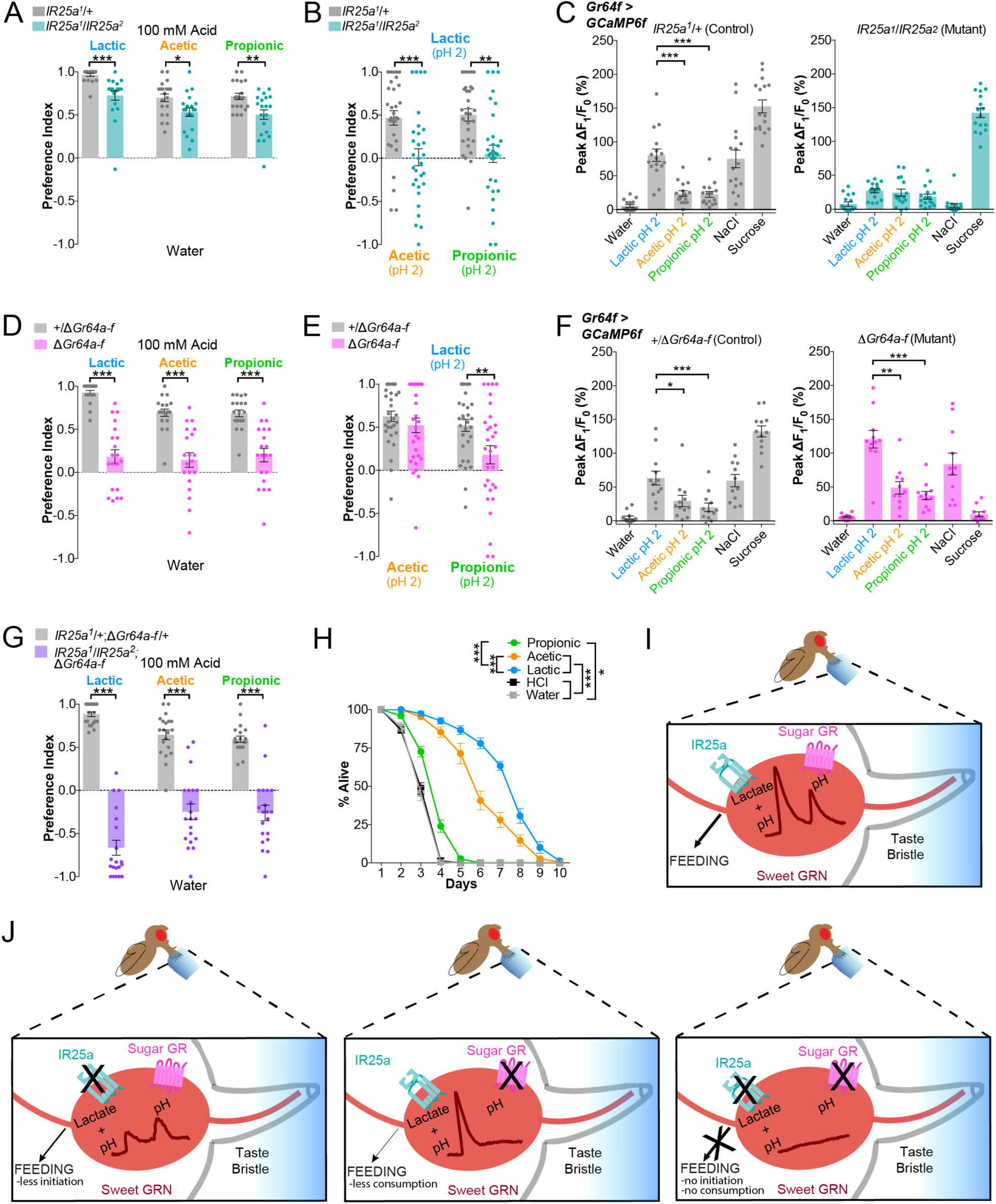
Differentiation between lactic and other attractive acids requires IR25a. (**A**) Binary choice feeding preference of *IR25a* mutants and heterozygous controls. Positive values indicate preference for 100 mM indicated acid; negative values indicate preference for water. Bars represent mean ±SEM. n=20 groups of 10 flies per genotype for each acid. Asterisks indicate significant differences by two-way ANOVA with Sidak’s post test, ∗p<.05, ∗∗p<.01, ∗∗∗p<.001. (**B**) Binary choice feeding of *IR25a* mutants and heterozygous controls in a volume-based assay with pH-matched solutions. Positive values indicate preference for 200 mM lactic acid pH=2; negative values indicate preference for 200 mM indicated acid (acetic or propionic) pH=2. Bars represent mean ±SEM. n=30 groups of 10 flies per genotype for each acid. Asterisks indicate significant differences by two-way ANOVA with Sidak’s post test, ∗∗p<.01, ∗∗∗p<.001. (**C**) Sweet GRN calcium responses in *IR25a* mutants and heterozygous controls with 4-second stimulations. Quantification of relative ‘onset’ peak fluorescence changes for each genotype. Bars represent mean ±SEM. n=16 flies per genotype. Asterisks indicate significant differences by one-way ANOVA with Dunnett’s post test, ∗∗∗p<.001. (**D**) Binary choice feeding preference of Δ*Gr64a-f* mutants and heterozygous controls. Positive values indicate preference for 100 mM indicated acid; negative values indicate preference for water. Bars represent mean ±SEM. n=20 groups of 10 flies per genotype for each acid. Asterisks indicate significant differences by two-way ANOVA with Sidak’s post test, ∗∗∗p<.001. (**E**) Binary choice feeding of Δ*Gr64a-f* mutants and heterozygous controls in a volume-based assay with pH-matched solutions. Positive values indicate preference for 200 mM lactic acid pH=2; negative values indicate preference for 200 mM indicated acid (acetic or propionic) pH=2. Bars represent mean ±SEM. n=29-31 groups of 10 flies per genotype for each acid. Asterisks indicate significant differences by two-way ANOVA with Sidak’s post test, ∗∗p<.01. (**F**) Sweet GRN calcium responses in Δ*Gr64a-f* mutants and heterozygous controls with 4-second stimulations. Quantification of relative ‘onset’ peak fluorescence changes for each genotype. Bars represent mean ±SEM. n=11-12 flies per genotype. Asterisks indicate significant differences by one-way ANOVA with Dunnett’s post test, ∗p<.05, ∗∗p<.01, ∗∗∗p<.001. (**G**) Binary choice feeding preference of *Ir25a +* Δ*Gr64a-f* mutants and heterozygous controls. Positive values indicate preference for 100 mM indicated acid; negative values indicate preference for water. Bars represent mean ±SEM. n=20 groups of 10 flies per genotype for each acid. Asterisks indicate significant differences by two-way ANOVA with Sidak’s post test, ∗∗∗p<.001. (**H**) Survival of control *w*^*1118*^ flies on water, 250 mM indicated acids, or HCl pH=2. Points represent mean ±SEM. n=15 groups of 10 flies per solution plotted as % of flies alive per group per day. Asterisks denote significance between groups by two-way ANOVA with Tukey’s post test, ∗p<.05, ∗∗∗p<.001. (**I**) Model of a single sweet GRN inside a taste bristle representing how sweet GRNs respond to lactic acid stimulation: the onset calcium peak is largely in response to lactate, which is specifically detected by IR25a, whereas the removal calcium peak is generated by a change in pH, which is mediated by sweet GRs. (**J**) Model summarizing the effects of eliminating each receptor type (individually or together) on the sweet GRN calcium kinetics and feeding behaviour.

Strikingly, control flies showed a clear preference for lactic acid over both other acids, but this preference was completely abolished in *IR25a* mutants (Fig. 7B). Moreover, calcium imaging of the onset peaks using these same, 200 mM pH-matched solutions, showed that lactic acid responses are significantly higher than the other acids in control flies, but all three acids produce equivalent responses in *IR25a* mutants (Fig. 7C).

Consistent with their behaviour towards lactic acid, Δ*Gr64a-f* mutants lose much of their attraction to acetic acid and propionic acid over water (Fig. 7D). However, in contrast to the results with *IR25a*, Δ*Gr64a-f* mutants retained strong preference for lactic acid over acetic acid in the pH-matched choice assay (Fig. 7E). Although there was a reduction in the preference for lactic acid over propionic acid in the mutants, the preference remained in the positive range towards lactic acid. Furthermore, calcium imaging of the onset peaks with these same solutions showed that lactic acid responses are significantly higher than the other acids in both controls and Δ*Gr64a-f* mutants (Fig. 7F). These results support the notion that sweet GRs are non-specifically responding to pH, and are less involved in the differentiation between attractive acids.

As with lactic acid, attraction to acetic acid and propionic acid was completely abolished in combined *IR25a*, Δ*Gr64a-f* mutants, leaving behavioural aversion (Fig. 7G). This suggests that all three acids are sensed by the same, or similar, mechanisms, but that the IR25a-containing receptor is more strongly tuned to lactate than to acetate or propionate.

Finally, to investigate a possible explanation for the relative behavioural attraction to other acids, we compared the ability of all three carboxylic acids to serve as a sole energy source. We found that lactic acid had the most significant positive impact on survival, followed closely by acetic acid (Fig. 7H). Propionic acid minimally prolonged survival, and HCl, as a low pH control, produced the same results as water alone.

## DISCUSSION

The receptors and channels involved in sour taste have been particularly difficult to identify because a large proportion of proteins have the potential to respond to acids either directly or indirectly (Holzer 2009). By using *Drosophila melanogaster* as a model organism, we were able to assess more nuanced aspects of acid detection *in vivo* and determine the impact of different acid components on feeding behaviour. Our results reveal an unprecedented complexity in the chemoreception of lactic acid, where different classes of receptors are required for the detection of the anion and pH and both are required for behavioural feeding attraction (Fig. 7I,J).

### Onset and removal responses in labellar GRNs

This is the first demonstration of sweet GRNs responding to both the onset and removal of a tastant, a phenomenon that was recently described for bitter GRNs (Devineni et al. 2020; Snell et al. 2020). In bitter GRNs, the same bitter GRs mediate both the onset and removal peaks to bitter compounds, and these distinct peaks are maintained in higher-order taste circuits to have meaningful physiological consequences (Devineni et al. 2020; Snell et al. 2020). While the full details of how the two sweet GRN peaks impact higher-order encoding of acids are unclear, our analysis suggests that both peaks positively contribute to feeding behaviour. Remarkably, the two peaks are mediated by distinct receptors, demonstrating that a single molecule is able to activate sweet GRNs through multiple mechanisms to ultimately produce behavioural attraction (Fig. 7I,J).

In *Drosophila*, there are two commonly used techniques to assess GRN activation by tastants: calcium imaging of molecularly-defined populations of neurons, and electrophysiological tip recordings of individual sensilla. While, historically, these two methods have largely led to similar conclusions, each has its own strengths. Of particular relevance here, calcium imaging allows for visualization of activity before, during, and after stimulation, which revealed the previously undescribed two-peak phenotype in sweet GRNs. This same methodology allowed for the recent description of onset and removal peaks in bitter GRNs (Devineni et al. 2020; Snell et al. 2020). By contrast, tip recordings measure activity only during stimulation, and recordings of L-type sensilla have not been found to respond to acid stimulation alone (Charlu et al. 2013; Rimal et al. 2019). Given that we observed heterogeneity in the areas of sweet GRN terminals activated during stimulus onset and removal, and a sugar-sensing neuron is located in each L, I, and S-type sensillum, it is possible that acids stimulate a subset of sugar GRNs outside of those measured in tip-recordings (Jaeger et al. 2018; Dahanukar et al. 2007). Future experiments can determine if different sensillum types differentially contribute to acid taste coding.

### A dual receptor mechanism in gustatory acid attraction

To our knowledge, this is the first instance where gustatory detection of a single compound requires two different receptor families, IRs and GRs, working in concert. IR25a appears to primarily mediate the onset peak in sweet GRNs, which correlates with feeding initiation (PER), and is likely driven by detection of the specific anion. Conversely, sweet GRs appear to dampen this onset peak. These GRs also mediate the sweet GRN removal peak, which is a non-specific response to acid removal and correlates with ingestion behaviour. One curious observation is that *IR25a* mutants retain a small onset peak and Δ*Gr64a-f* mutants have an enhanced onset, yet the combined mutants show no response at all. We speculate that GRs and IR25a both respond to onset of low pH to some extent, and in the absence of GRs, enhancement of the IR25a-dependent pH response masks loss of the small GR-dependent onset response. While this is technically challenging to resolve, future experiments can attempt to do so.

We were surprised that IR76b did not contribute to lactic acid attraction, given its overlapping function with IR25a in many other instances of chemoreception (Jaeger et al. 2018; Lee et al. 2018; Ahn, Chen, and Amrein 2017; Y. Chen and Amrein 2017; Sánchez-Alcañiz et al. 2018). Instead, it appears to be involved in limiting lactic acid attraction. Both IRs are expressed broadly across many classes of GRNs on the labellum (Lee et al. 2018; Jaeger et al. 2018) and we cannot rule out that they are contributing to the detection of acids in bitter GRNs. However, the results with *IR76b* mutants fits with a proposed role for IR76b in limiting sensitivity directly in sweet GRNs (H.-L. Chen, Stern, and Yang 2019).

The involvement of sweet GRs in acid attraction was also surprising. *Drosophila melanogaster* has nine known sugar GRs which are well characterized in their detection of specific sugars (Yavuz et al. 2014; Dahanukar et al. 2007; Jiao et al. 2008; Slone, Daniels, and Amrein 2007; Miyamoto et al. 2012; Dahanukar et al. 2001; Wisotsky et al. 2011). Similar to IRs, these GRs are thought to function as multimers, with Gr64f being a possible co-receptor for sugar detection (Jiao et al. 2008). Recently, Gr64e, which acts as a glycerol receptor, was also shown to have a non-canonical role in fatty acid taste transduction downstream of phospholipase C (PLC) (Kim et al. 2018). This demonstrates that a single GR can act as both a ligand-gated ion channel in the direct reception of sweet compounds and indirectly contribute to the detection of other molecules. However, acetic acid taste was previously shown to be independent of the PLC pathway, suggesting that this is also not the function of GRs in lactic acid taste (Devineni et al., 2019). Our data suggests that all nine sugar GRs contribute to lactic acid feeding. It is unclear how lactic acid taste is so sensitive to GR dose, but this is a likely explanation for why single GR mutants show partial phenotypes in the binary choice feeding assay. The function of GRs in acid taste appears to be a non-specific response to pH changes. This is consistent with the acid sensitivity of bitter GRNs, which also express a large complement of GRs. We speculate that in sweet GRNs at rest, GRs are in a configuration that either limits the amount of acid entering the cell or limits the response of IR25a to low pH. With sufficient acidification of the lymph and/or intracellular fluid, the gating or conformation of GRs may change so that relief from acidity results in additional ion flux.

### Lactic acid as a particularly attractive sensory stimulus

Our results confirm that lactic acid is particularly attractive to *Drosophila*, more so than other carboxylic acids commonly used in behavioural chemosensory experiments. Surprisingly, lactic acid has not been used in *Drosophila* olfaction studies, despite being a key olfactory attractant to mosquitoes (McBride 2016). We found that lactic acid smell is attractive to *Drosophila* and that the olfactory co-receptor IR8a is required for this attraction, similar to mosquitoes (Raji et al. 2019). Previously, IR8a was found to mediate aversion to acetic acid in IR64+ olfactory neurons in *Drosophila*; however, IR8a is located in additional IR64-glomeruli (Ai et al. 2013). Lactic acid likely activates both attractive and aversive olfactory neurons, similar to other acidic stimuli (Semmelhack and Wang 2009). Removing olfactory organs completely had no impact on lactic acid feeding attraction, whereas *IR8a* mutation alone led to a small but significant reduction in feeding preference. These results highlight the complexity of the full chemosensory response to lactic acid when factoring in olfactory pathways in addition to gustation. Future experiments can further explore this olfactory mechanism of lactic acid attraction in *Drosophila*, and conversely, explore how lactic acid gustatory sensing in mosquitoes may influence biting.

We briefly explored one potential reason behind the strong attraction to lactic acid in *Drosophila*, by investigating its ability to provide energy in nutrient-deprived flies. Lactic acid was particularly effective in improving fly survival, likely because lactate is a fuel for the TCA cycle (Rabinowitz and Enerbäck 2020). Acetic acid as a potential source of energy was speculated to be one reason for attraction in a previous *Drosophila* study (Devineni et al. 2019), and we find that acetic acid provides energy to flies, but less efficiently than lactic acid.

Propionic acid only minimally increased survival, and the more limited metabolic pathways associated with these other acids could explain their less efficient function as alternative energy sources. Additional explanations, such as attraction to specific gut microbes, undoubtedly exist, but the use of carboxylic acids as fuel provides one explanation for feeding attraction.

## ACKNOWLEDGEMENTS

We thank Anupama Dahanukar, the Bloomington Stock Center, and the Vienna Drosophila Resource Center for fly stocks, and members of the Gordon lab for comments on the manuscript. This work was funded by the Canadian Institutes of Health Research (CIHR) operating grant FDN-148424. M.D.G. is a Michael Smith Foundation for Health Research Scholar.

## AUTHOR CONTRIBUTIONS

M.S. and M.D.G conceived the project and wrote the manuscript. M.S. performed all calcium imaging, survival experiments, PER assays, CAFE assays, trap assays, immunostaining, data analysis, and binary choice experiments not listed below. B.G. performed binary choice assays for candidate receptors, the GRN silencing screen, other acid attraction in control flies, and for flies with surgical removal of olfactory organs. Z.F.W contributed to the initial binary choice assays in control flies. J.C. contributed to the binary choice GRN silencing screen. M.D.G. supervised the project.

## DECLARATION OF INTERESTS

The authors declare no competing interests.

## SUPPLMENTAL INFORMATION

Supplemental information includes six figures.

## MATERIALS AND METHODS

### Flies

Flies were raised on standard cornmeal fly food at 25°C in 70% humidity. All experiments were performed on 2-10 day-old flies using mated females unless stated otherwise. Genotypes used in each experiment are listed below, additional source and strain information can be found in the Key Resources Table.

Figure 1:

- *w1118*
- *Ir8a*^*1*^*/+; +/+; +/+*
- *Ir8a*^*1*^*/Ir8a*^*1*^*;+/+;+/+*

Figure 2:

- +/+; +/+; *UAS-Kir2*.*1,tub-Gal80*^*TS*^/+
- +/+; *Gr64f-Gal4*/+; +/+
- +/+; *Gr64f-Gal4*/+; *UAS-Kir2*.*1,tub-Gal80*^*TS*^/+
- +/+; +/+; *Ir94e-Gal4*/+
- +/+; +/+; *Ir94e-Gal4/UAS-Kir2*.*1,tubGal80*^*TS*^
- *Gr66a-LexA/+; LexAop-Gal80*/+; *Ppk23-Gal4*/+
- *Gr66a-LexA*/+; *LexAop-Gal80*/+; *Ppk23-Gal4/UAS-Kir2*.*1,tub-Gal80*^*TS*^
- +/+; +/+; *Ppk28-Gal4*/+
- +/+; +/+; *Ppk28-Gal4/UAS-Kir2*.*1,tub-Gal80*^*TS*^
- +/+; *Gr66a-Gal4*/+; +/+
- +/+; *Gr66a-Gal4/+; UAS-Kir2*.*1,tub-Gal80*^*TS*^/+
- +/+; *Gr64f-Gal4/UAS-GCaMP6f*; /+
- +/+; *IR56d-Gal4*/*UAS-GCaMP6f*; +/+

Figure 3:

- +/+; *Gr64f-Gal4/UAS-GCaMP6f*; /+

Figure 4:

- +/+; *IR25a*^*1*^/+; +/+
- +/+; *IR25a*^*2*^/+; +/+
- +/+; *IR25a*^*1*^/*IR25a*^*2*^; +/+
- +/+; *IR25a*^*1*^/+;*UAS-IR25a*/+
- +/+; I*R25a*^*2*^/+; *Gr64f-Gal4*/+
- +/+; *IR25a*^*1*^*/IR25a*^*2*^; *UAS-IR25a*/+
- +/+; I*R25a*^*1*^/*IR25a*^*2*^; *Gr64f-Gal4*/+
- +/+; *IR25a*^*1*^/*IR25a*^*2*^; *Gr64f-Gal4*/*UAS-IR25a*
- +/+; +/+; *IR76b*^*1*^/+
- +/+; +/+; *IR76b*^*2*^/+
- +/+; +/+; *IR76b*^*1*^/*IR76b*^*2*^
- +/+; *IR25a*^*1*^/*+*; *Gr64f-Gal4*/*UAS-GCaMP6f*
- +/+; *IR25a*^*1*^/*IR25a*^*2*^; *Gr64f-Gal4*/*UAS-GCaMP6f*

Figure 5:

- Δ*Gr5a*/+; +/+; +/+
- Δ*Gr5a*/ Δ*Gr5a*; +/+; +/+
- +/+; *Gr43a*^*LEXA*^/+; +/+
- +/+; *Gr43a*^*GAL4*^/*Gr43a*^*GAL4*^; +/+
- +/+; Δ*Gr61a*^*1*^/+; +/+
- +/+; Δ*Gr61a*^*1*^/Δ*Gr61a*^*1*^; +/+
- +/+; Δ*Gr64a*^*1*^/+; +/+
- +/+; Δ*Gr64a*^*1*^/Δ*Gr64a*^*1*^; +/+
- +/+; Δ*Gr64a*^*2*^/+; +/+
- +/+; Δ*Gr64a*^*2*^/Δ*Gr64a*^*2*^; +/+
- +/+; Δ*Gr64d*^*1*^/+; +/+
- +/+; Δ*Gr64d*^*1*^/Δ*Gr64d*^*1*^; +/+
- +/+; Δ*Gr64e*^*MB03533*^/+; +/+
- +/+; Δ*Gr64e*^*MB03533*^/Δ*Gr64e*^*MB03533*^; +/+
- +/+; +/+; *Gr64f*^*LEXA*^/+
- +; +/+; *Gr64f*^*LEXA*^/*Gr64f*^*LEXA*^
- +/+; +/+; Δ*Gr64a-f*/+
- +/+; +/+; Δ*Gr64a-f*/Δ*Gr64a-f*
- Δ*Gr5a*/+; Δ*Gr64a*/+; +/+
- Δ*Gr5a*/Δ*Gr5a*; Δ*Gr64a*/Δ*Gr64a*; +/+
- *R1,Gr5a*^*LEXA*^/+; +/+; Δ*Gr61a*, Δ*Gr64a-f*/+
- *R1,Gr5a*^*LEXA*^/*R1,Gr5a*^*LEXA*^; +/+; Δ*Gr61a*, Δ*Gr64a-f*/Δ*Gr61a*, Δ*Gr64a-f*
- *R1,Gr5a*^*LEXA*^/+; *Gr43a*^*LEXA*^/+; Δ*Gr61a*, Δ*Gr64a-f*/+
- *R1,Gr5a*^*LEXA*^/*R1,Gr5a*^*LEXA*^; *Gr43a*^*LEXA*^/*Gr43a*^*LEXA*^; Δ*Gr61a*, Δ*Gr64a-f*/Δ*Gr61a*, Δ*Gr64a-f*
- +/+; *Gr64f-Gal4,UAS-GCaMP6f*/+; Δ*Gr64a-f*/+
- +/+; *Gr64f-Gal4,UAS-GCaMP6f*/+; Δ*Gr64a-f*/Δ*Gr64a-f*

Figure 6:

- +/+; *IR25a*^*1*^,*Gr64f-Gal4*/*UAS-GCaMP6f*; Δ*Gr64a-f*/+
- +/+; *IR25a*^*1*^,*Gr64f-Gal4*/*IR25a*^*2*^,*UAS-GCaMP6f*; Δ*Gr64a-f*/Δ*Gr64a-f*
- +/+; *IR25a*^*1*^/+; Δ*Gr64a-f*/+
- +/+; *IR25a*^*1*^/*IR25a*^*2*^; Δ*Gr64a-f*/+
- +/+; *IR25a*^*1*^/+; Δ*Gr64a-f*/Δ*Gr64a-f*
- +/+; *IR25a*^*1*^/*IR25a*^*2*^; Δ*Gr64a-f*/Δ*Gr64a-f*

Figure 7:

- +/+; *IR25a*^*1*^/+; +/+
- +/+; *IR25a*^*1*^/*IR25a*^*2*^; +/+
- +/+; *IR25a*^*1*^/*+*; *Gr64f-Gal4*/*UAS-GCaMP6f*
- +/+; *IR25a*^*1*^/*IR25a*^*2*^; *Gr64f-Gal4*/*UAS-GCaMP6f*
- +/+; +/+; Δ*Gr64a-f*/+
- +/+; +/+; Δ*Gr64a-f*/Δ*Gr64a-f*
- +/+; *Gr64f-Gal4,UAS-GCaMP6f*/+; Δ*Gr64a-f*/+
- +/+; *Gr64f-Gal4,UAS-GCaMP6f*/+; Δ*Gr64a-f*/Δ*Gr64a-f*
- +/+; *IR25a*^*1*^/+; Δ*Gr64a-f*/+
- +/+; *IR25a*^*1*^/*IR25a*^*2*^; Δ*Gr64a-f*/Δ*Gr64a-f*

Figure S1:

- *w*^*1118*^
- *Canton S*.
- *Ir8a*^*1*^*/+; +/+; +/+*
- *Ir8a*^*1*^*/Ir8a*^*1*^*;+/+;+/+*

Figure S2:

- +/+; *Gr64f-Gal4*/+; *UAS-mCD8::GFP*/+
- +/+; +/+; *UAS-Kir2*.*1,tub-Gal80*^*TS*^/+
- +/+; *Gr64f-Gal4*/+; +/+
- +/+; *Gr64f-Gal4*/+; *UAS-Kir2*.*1,tub-Gal80*^*TS*^/+
- +/+; *Gr64e-Gal4*/+; *UAS-mCD8::GFP*/+
- +/+; +/+; *UAS-Kir2*.*1,tub-Gal80*^*TS*^/+
- +/+; *Gr64e-Gal4*/+; +/+
- +/+; *Gr64e-Gal4*/+; *UAS-Kir2*.*1*/+

Figure S3:

- +/+; *Gr66a-Gal4*/*UAS-GCaMP6f*; +/+

Figure S4:

- +/+; *IR25a*^*1*^/+; *Gr64f-Gal4*/*UAS-GCaMP6f*
- +/+; *IR25a*^*1*^/*IR25a*^*2*^; *Gr64f-Gal4*/*UAS-GCaMP6f*
- +/+; *IR25a*^*1*^,*UAS-IR25a*/*IR25a*^*2*^; *Gr64f-Gal4*/*UAS-GCaMP6f*

Figure S5:

- +/+; *Gr64f-Gal4*/+; +/+
- +/+; *Gr64f-Gal4*/*UAS-Ir7c RNAi*; +/+
- +/+; *Gr64f-Gal4*/+; *UAS-IR56d RNAi*/+
- +/+; *IR56b*^*GAL4*^+; +/+
- +/+; *IR56b*^*GAL4*^/*IR56b*^*GAL4*^; +/+
- +/+; +/+; Δ*IR62a*/Δ*IR62a*
- +/+; *Gr64f-Gal4,UAS-GCaMP6f*/+; Δ*Gr64a-f*/+
- +/+; *Gr64f-Gal4,UAS-GCaMP6f*/+; Δ*Gr64a-f*/Δ*Gr64a-f*

Figure S6:

- *w*^*1118*^
- +/+; *Gr64f-Gal4/UAS-GCaMP6f*; /+

### Tastants

The following tastants were used: DL-lactic acid, sucrose, NaCl, caffeine, acetic acid, propionic acid, hydrochloric acid (Sigma-Aldrich). Tastants were kept as 1 M stocks and diluted as necessary for experiments. The pH of tastants were adjusted where indicated using concentrated HCl or NaOH.

### Behavioural assays

Olfactory trap assays were designed to resemble previous protocols (Ogueta et al. 2010; Stensmyr et al. 2012). Groups of 40 flies were starved on 1% agar for 2 hours prior to the assay. Flies were lightly anesthetized with CO_2_ and placed in the trap assay which consisted of a glass container (11 cm diameter x 11 cm height) containing two 25 mL glass flasks with 10 mL of either ddH_2_O or 250 mM lactic acid. The flasks were sealed with parafilm except for a small hole in the middle where a 1000mL pipette tip was placed, stopping ∼2 cm from the top of the solutions. The top of the tip was cut to ∼8 mm and bottom of the tip was cut to ∼2.5 mm, and the parafilm made contact with the pipette tip so that there were no potential exits from the flasks.

The lid of the glass container had mesh holes larger than the flies, so parafilm was used to cover the mesh and 100 small holes poked uniformly throughout for airflow. Flies in the trap assay were placed at 29°C in the dark for ∼18 hrs. After, flies were anesthetized with CO_2_ and the number of flies choosing the flask with water, lactic acid, or neither were counted and a preference index (PI) calculated as: ((# of flies in lactic acid flask)-(# of flies in water flask))/(total # of flies in either flask).

Binary choice feeding assays were performed similarly to previous descriptions (Jaeger et al. 2018). Groups of 10 flies were starved on 1% agar for 1 day at 25°C prior to testing. For neural silencing with Kir2.1, expression was induced by placing flies at 29°C for three days prior to experiment to inactivate Gal80^ts^ (2 days on food and 1 day on 1% agar). For all binary choice experiments, flies were transferred into vials containing six 10 μL drops of alternating color.

Each drop contained the specified concentration of tastant in 1% agar with either blue (0.125mg/mL Erioglaucine, FD and C Blue#1) or red (0.5mg/mL Amaranth, FD and C Red#2) dye. Color was balanced for each experiment (i.e. half of the replicates had lactic acid in red, water in blue, and half of the replicates had water in red, lactic acid in blue). Flies were allowed to feed for 2 hrs at 29°C in the dark before freezing at -20°C. Abdomen color was scored under a dissection microscope as red, blue, purple, or no color. PI was calculated as ((# of flies labeled with tastant 1 color)-(# of flies labeled with tastant 2 color))/(total # of flies with color). Any vials with <30% of flies feeding were excluded.

Capillary Feeder (CAFE) assays quantified over 24 hrs (Fig. 1) were performed as previously described (Stafford et al. 2012). Briefly, 10 flies were starved for 5 hours and then placed in specialized 15 ml conical vials with access to two capillary tubes (A-M Systems 626000) containing water or two capillary tubes containing 250 mM lactic acid. All solutions contained 0.01% FD&C Blue No. 1 dye for visualization in photographs of the capillaries, which were taken once per hour for 24 hrs with a Pentax Optio W90 handheld digital camera at 29°C. Two vials in each experiment did not contain flies and were used to control for the volume change due to evaporation. ImageJ was used to calculate the volume of solution consumed in each capillary from the photographs. PI was calculated by ((volume consumed of lactic acid)-(volume consumed of water)/(total volume consumed)) for each 4-hr interval over 24 hrs. An acute CAFE binary assay was used in feeding experiments using pH-matched solutions (Fig. 6). For these experiments, flies were starved 24 hrs prior to the start and the same CAFE protocol was used except no dye was added to the solutions. The volume was marked by hand on the capillary tubes at the start and after 4 hrs of feeding at 29°C. The distance between marks (i.e. volume consumed) was quantified in mm using a standard ruler under a dissection microscope. PI was calculated the same as above.

PER was performed as previously described (Jaeger et al. 2018; Stafford et al. 2012). For labellar PER, flies were mounted inside of 200 μL pipette tips cut so that only the heads were exposed.

Tubes were sealed with tape on the bottom and placed onto a slide with double sided tape. For tarsal PER, flies were immobilized on slides containing strips of myristic acid. For both assays, after a 1-2 hr recovery in a humidity chamber, flies were stimulated with water and allowed to drink until satiated (flies showing continued extension to water were excluded). Each fly was stimulated on either the labellum or tarsi with increasing concentrations of lactic acid followed by 500 mM sucrose as a positive control using a 20 μL pipette attached to a 1 mL syringe. Each tastant was presented one time and water was offered in between each tastant to maintain satiation. For each tastant, flies showing clear extension were scored as 1 for that tastant, or 0 if not, and data are plotted as percent responding. Experiments were conducted over four different days, using 10-15 flies per genotype matched each day, and the order of genotypes stimulated each day was randomized.

For behavioural experiments with no olfactory organs, experimental flies were anesthetized with CO_2_ after collection and sharp forceps used to remove the 3^rd^ segment of the antennae and the maxillary palps of each fly. Control flies were anesthetized with CO_2_ for the same duration. Both groups were allowed to recover for ∼2 hrs before being moved to starvation vials prior to the start of the experiments.

### Calcium imaging

*In vivo* GCaMP imaging of GRN axon terminals was performed as previously described (Jaeger et al. 2018). Mated female flies aged 2-6 days were briefly anesthetized with CO_2_ and placed in a custom chamber. Nail polish was used to secure the back of the neck and a small amount of wax was applied to both sides of the proboscis in an extended position, covering the maxillary palps without touching the labellar sensilla. After 1 hr recovery in a humidity chamber, antennae were removed along with a small window of cuticle to expose the SEZ. Adult hemolymph-like (AHL) solution (108 mM NaCl, 5 mM KCl, 4 mM NaHCO3, 1 mM NaH2PO4, 5 mM HEPES, 15 mM ribose, 2mM Ca^2+^, 8.2mM Mg^2+^, pH 7.5) was immediately applied. Air sacs and fat were removed and the esophagus was clipped and removed for clear visualization of the SEZ.

A Leica SP5 II Confocal microscope was used to capture GCaMP6f fluorescence with a 25x water immersion objective. The SEZ was imaged at a zoom of 4x, line speed of 8000 Hz, line accumulation of 2, and resolution of 512 × 512 pixels. Pinhole was opened to 2.86 AU. For each taste stimulation, 15 total seconds were recorded. For 1 s stimulations, this consisted of 5 s baseline, 1 s stimulation, 9 s post-stimulation. For 4 s stimulations, this consisted of 5 s baseline, 4 s stimulation, 6 s post-stimulation. A pulled capillary filed down to fit over both labellar palps was filled with tastant and positioned close to the labellum with a micromanipulator. For the stimulation, the micromanipulator was manually moved over the labellum and then removed from the labellum after 1 or 4 s. The stimulator was washed with water in between tastants of differing solutions.

The maximum change in fluorescence (peak ΔF/F) for ON peaks was calculated using peak intensity (average of 3 time points) minus the average baseline intensity (10 time points), divided by the baseline. For OFF peaks, peak ΔF/F was calculated using peak intensity during stimulus removal (average of 3 time points) minus the minimum intensity prior to removal (3 time points), divided by the minimum intensity (a new ‘baseline’). ImageJ was used to quantify fluorescence changes and create heatmaps.

### Survival experiments

Adult, mated female flies, were collected and placed on the indicated solution as the only food option in 1% agar at room temperature at 3 days old. Flies were flipped onto fresh solution in agar every two days. A total of ten flies were in each vial and the number dead and alive counted once per day, and plotted as the % of flies alive. All solutions were run in parallel.

### Immunohistochemistry

Brain immunofluorescence was performed as previously described (Jaeger et al. 2018). Primary antibodies used were rabbit anti-GFP (1:1000, Invitrogen) and mouse anti-brp (1:50, DSHB #nc82). Secondary antibodies used were goat anti-rabbit Alexa 488 and goat anti-mouse Alexa 546 (1:200, Invitrogen). Images were acquired using a Leica SP5 II Confocal microscope under 25x objective. Images were processed in ImageJ.

### Statistical analysis

Statistical tests were performed using GraphPad Prism 6 software and are stated in the figure legends along with the sample sizes and what is considered a biological replicate for each experiment. Sample sizes were generally determined prior based on the variance and effect sizes seen in previous experiments. Experimental conditions and genotype controls were run in parallel. In the rare occurrence that a data point visibly appeared to be an outlier, a Grubb’s test was performed and excluded from the data set if meeting a significance value of <.05. In the calcium imaging experiments using *Gr64a-f* mutants, ∼15% showed a significant response to water (with stimulus onset or removal) and were removed from the final dataset.

## “Key Resources Table”

| **Reagent type (species) or resource** | **Designation** | **Source or reference** | **Identifiers** |
| --- | --- | --- | --- |
| Genetic reagent (*D. melanogaster*) | *w<sup>1118</sup>* | Bloomington Drosophila Stock Center | BDSC: 32214 |
| Genetic reagent (*D. melanogaster*) | *Canton S.* | Bloomington Drosophila Stock Center | BDSC: 64349 |
| Genetic reagent (*D. melanogaster*) | *IR8a<sup>1</sup>* | Bloomington Drosophila Stock Center | BDSC: 41744 |
| Genetic reagent (*D. melanogaster*) | *UAS-Kir2.1* | (Baines et al. 2001) | Flybase: FBti0017552 |
| Genetic reagent (*D. melanogaster*) | *Tub-Gal80<sup>TS</sup>* | (McGuire, Mao, and Davis 2004) | Flybase: FBti0027797 |
| Genetic reagent (*D. melanogaster*) | *Ir94e-Gal4* | (Tirian and Dickson 2017; Jaeger et al. 2018) | VDRC: v207582 |
| Genetic reagent (*D. melanogaster*) | *Gr66a-LexA* | (Thistle et al. 2012) | Flybase: FBal0277069 |
| Genetic reagent (*D. melanogaster*) | *LexAop-Gal80* | Bloomington Drosophila Stock Center | BDSC: 44277 |
| Genetic reagent (*D. melanogaster*) | *Ppk28-Gal4* | (Cameron et al. 2010) | Flybase: FBtp0054514 |
| Genetic reagent (*D. melanogaster*) | *Gr66a-Gal4* | (Z. Wang et al. 2004) | Flybase: FBtp0014660 |
| Genetic reagent (*D. melanogaster*) | *Gr64f-Gal4* | (Dahanukar et al. 2007) | Flybase: FBti0162678 |
| Genetic reagent (*D. melanogaster*) | *Gr64f-Gal4* | (Dahanukar et al. 2007) | Flybase: FBtp0057275 |
| Genetic reagent (*D. melanogaster*) | *IR56d-Gal4* | (Sánchez-Alcañiz et al. 2018) | BDSC: 81235 |
| Genetic reagent (*D. melanogaster*) | *UAS-GCaMP6f* | Bloomington Drosophila Stock Center | BDSC: 42747; |
| Genetic reagent (*D. melanogaster*) | *UAS-GCaMP6f* | Bloomington Drosophila Stock Center | BDSC: 52869; |
| Genetic reagent (*D. melanogaster*) | *IR25a<sup>1</sup>* | (Benton et al. 2009) | Flybase: FBst0041736 |
| Genetic reagent (*D. melanogaster*) | *IR25a<sup>2</sup>* | (Benton et al. 2009) | Flybase: FBst0041737 |
| Genetic reagent (*D. melanogaster*) | *UAS-IR25a* | Bloomington Drosophila Stock Center | BDSC: 78067 |
| Genetic reagent (*D. melanogaster*) | *IR76b<sup>1</sup>* | (Y. V. Zhang, Ni, and Montell 2013) | Flybase: FBst0051309 |
| Genetic reagent (*D. melanogaster*) | *IR76b<sup>2</sup>* | (Y. V. Zhang, Ni, and Montell 2013) | Flybase: FBst0051310 |
| Genetic reagent (*D. melanogaster*) | *ΔIR62a* | Bloomington Drosophila Stock Center | BDSC: 32713 |
| Genetic reagent (*D. melanogaster*) | *IR56b<sup>GAL4</sup>* | Bloomington Drosophila Stock Center | BDSC: 27818 |
| Genetic reagent (*D. melanogaster*) | *UAS-IR56d RNAi* | Vienna Drosophila Resource Center | VDRC: 6112 |
| Genetic reagent (*D. melanogaster*) | *UAS-IR7c RNAi* | Vienna Drosophila Resource Center | VDRC: 109485 |
| Genetic reagent (*D. melanogaster*) | *ΔGr5a* | (Dahanukar et al. 2001) | Flybase: FBal0127256 |
| Genetic reagent (*D. melanogaster*) | *Gr43a<sup>GAL4</sup>* | (Miyamoto et al. 2012) | Flybase: FBal0290232 |
| Genetic reagent (*D. melanogaster*) | *ΔGr61a<sup>1</sup>* | (Dahanukar et al. 2007) | Flybase: FBal0256895 |
| Genetic reagent (*D. melanogaster*) | *ΔGr64a<sup>1</sup>* | (Dahanukar et al. 2007) | Flybase: FBal0256892 |
| Genetic reagent (*D. melanogaster*) | *ΔGr64a<sup>2</sup>* | (Dahanukar et al. 2007) | Flybase: FBab0047074 |
| Genetic reagent (*D. melanogaster*) | *ΔGr64d<sup>1</sup>* | (Uchizono et al. 2017) | Flybase: FBal0346605 |
| Genetic reagent (*D. melanogaster*) | *ΔGr64e<sup>MB03533</sup>* | (Wisotsky et al. 2011) | Flybase: FBal0192448 |
| Genetic reagent (*D. melanogaster*) | *Gr64f<sup>LEXA</sup>* | (Yavuz et al. 2014) | Flybase: FBal0304291 |
| Genetic reagent (*D. melanogaster*) | *ΔGr64a-f* | (Kim et al. 2018) | Flybase: FBab0049044 |
| Genetic reagent (*D. melanogaster*) | *R1,Gr5a<sup>LEXA</sup>; ; ΔGr61a, ΔGr64a-f* | (Yavuz et al. 2014) | Flybase: FBal0304286<br>FBal0256895<br>FBab0047080 |
| Genetic reagent (*D. melanogaster*) | *Gr64e-Gal4* | (Kwon et al. 2011) | BDSC: 57666 |
| Genetic reagent (*D. melanogaster*) | *UAS-mCD8::GFP* | Bloomington Drosophila Stock Center | BDSC: 32195 |
| Chemical compound, drug | DL-Lactic Acid | Sigma-Aldrich | 69785 |
| Chemical compound, drug | Sucrose | Sigma-Aldrich | S7903 |
| Chemical compound, drug | NaCl | Sigma-Aldrich | S7653 |
| Chemical compound,<br>drug | Caffeine | Sigma-Aldrich | C0750 |
| Chemical compound,<br>drug | NaOH | Sigma-Aldrich | 1310-23-10 |
| Chemical compound,<br>drug | Acetic Acid | Sigma-Aldrich | 64-19-7 |
| Chemical compound,<br>drug | Propionic Acid | Sigma-Aldrich | 79-09-4 |
| Chemical compound,<br>drug | Hydrochloric Acid | Sigma-Aldrich | 351280-212 |
| Chemical compound,<br>drug | Agar | Sigma-Aldrich | A1296 |
| Chemical compound,<br>drug | Erioglaucine, FD&C<br>Blue #1 | Spectrum | FD110 |
| Chemical compound,<br>drug | Amaranth FD&C Red<br>#2 | Sigma-Aldrich | A1016 |
| Chemical compound,<br>drug | 4% Paraformaldehyde<br>in PBS | Alfa Aesar | J61899 |
| Antibody | Mouse anti-brp | DSHB | Nc82 |
| Antibody | Rabbit anti-GFP | Invitrogen | A11122 |
| Antibody | Anti-rabbit Alexa 488 | Invitrogen | A11008 |
| Antibody | Anti-mouse Alexa 546 | Invitrogen | A11030 |
| Software | ImageJ | (Schneider, Rasband, and<br>Eliceiri 2012) | RRID: SCR_003070 |
| Software | Prism 6 | Graphpad | RRID: SCR_002798 |
| Software | Illustrator | Adobe | RRID: SCR_010279 |

## Notes

### Competing Interest Statement

The authors have declared no competing interest.

