## Supplemental information for "Mechanisms of lactic acid gustatory attraction in *Drosophila*"

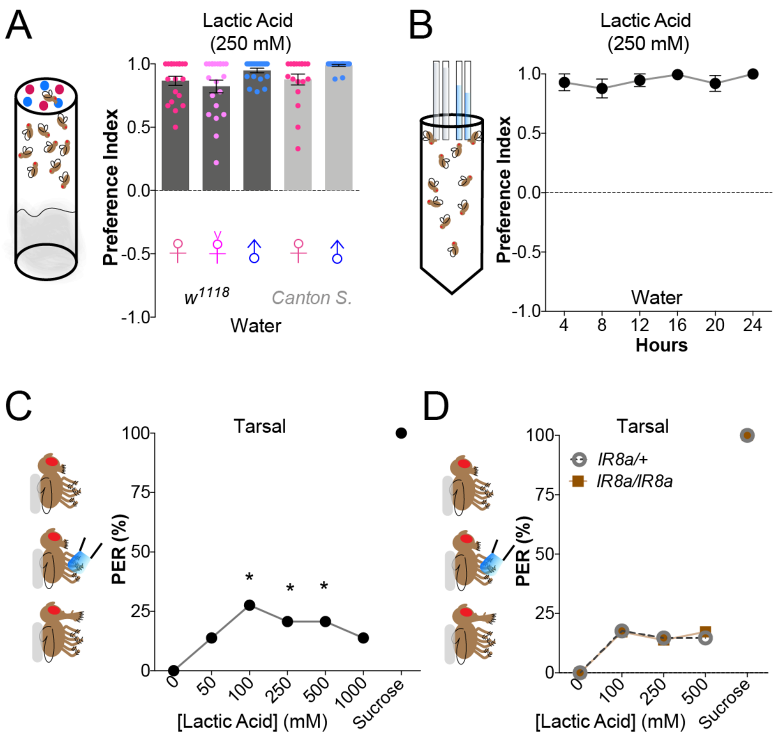


**Figure S1, related to Figure 1:**
**Lactic acid is broadly attractive to *Drosophila* despite minimal tarsal PER**
(**A**) Preferences of mated *w^1118^* females, virgin *w^1118^* females, *w^1118^* males, and wildtype *Canton S.* females and males in the binary feeding assay. Positive values indicate preference for 250 mM lactic acid; negative values indicate preference for water. No significant differences by one-way ANOVA with Dunnett's post test compared to mated *w^1118^* females. Bars represent mean ±SEM. n=20 groups of 10 flies per group. (**B**) Schematic of the volume-based capillary feeding assay (left) and lactic acid attraction in control *w^1118^* flies using this assay (right). Positive values indicate preference for 250 mM lactic acid; negative values indicate preference for water. Each point represents mean ±SEM for 7 vials of 10 flies each, with repeated volume and preference calculations at each time point per vial. (**C**) Tarsal PER of control *w^1118^* flies. n=29 flies and dots represent the mean. Each fly either responds by extending proboscis (1) or not (0) for each tastant presented in order, plotted as % responding. 500 mM sucrose was used as a positive control. Asterisks denote significant difference from negative control (0 mM, water) by one-way ANOVA with Dunnett's post test, *p<.05. (**D**) Tarsal PER of *IR8a* mutants and heterozygous controls. n=29-34 flies per genotype and dots represent the mean. 500 mM sucrose was used as a positive control. No significant differences by two-way ANOVA with Tukey's post test.

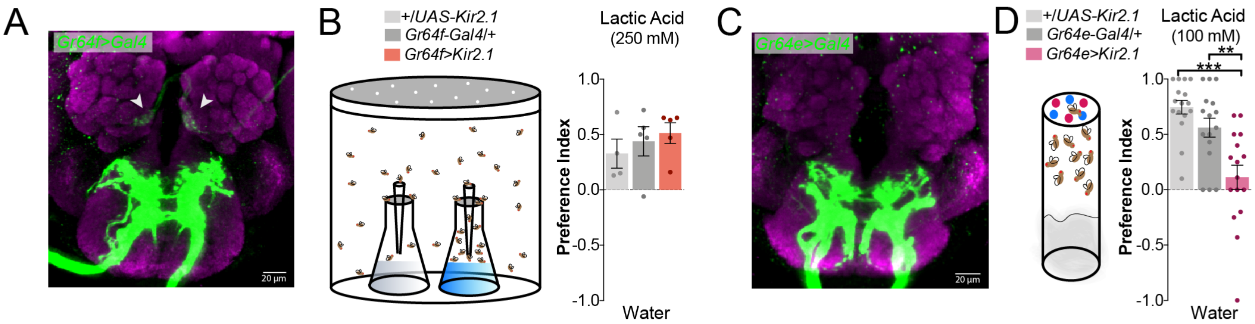


**Figure S2, related to Figure 2:**
**Sweet GRN silencing phenotypes are not driven by olfactory expression**
(**A**) Immunostained brains showing expression of *Gr64f-Gal4* driving *UAS-GFP* (green) and neuropil stained with nc82 (pink). Arrows indicate expression in antennal lobe coming from ORN projections in *Gr64f-Gal4*. (**B**) Olfactory attraction of flies with sweet GRNs silenced using *Gr64f-Gal4*, and genotype controls, in the trap assay. Positive values indicate preference for 250 mM lactic acid; negative values indicate preference for water. Bars represent mean ±SEM. n=4-5 sets of 40 flies per genotype. No significant differences by one-way ANOVA with Tukey's post test. (**C**) Immunostained brains showing expression of *Gr64e-Gal4* driving *UAS-GFP* (green) and neuropil stained with nc82 (pink). (**D**) Preferences in the binary feeding assay following sweet GRNs silencing with *Gr64e-Gal4*. Positive values indicate preference for 100 mM lactic acid; negative values indicate preference for water. Bars represent mean ±SEM. n=16 groups of 10 flies per genotype. Asterisks denote significance between by one-way ANOVA with Tukey's post test, **p<.01, ***p<.001.

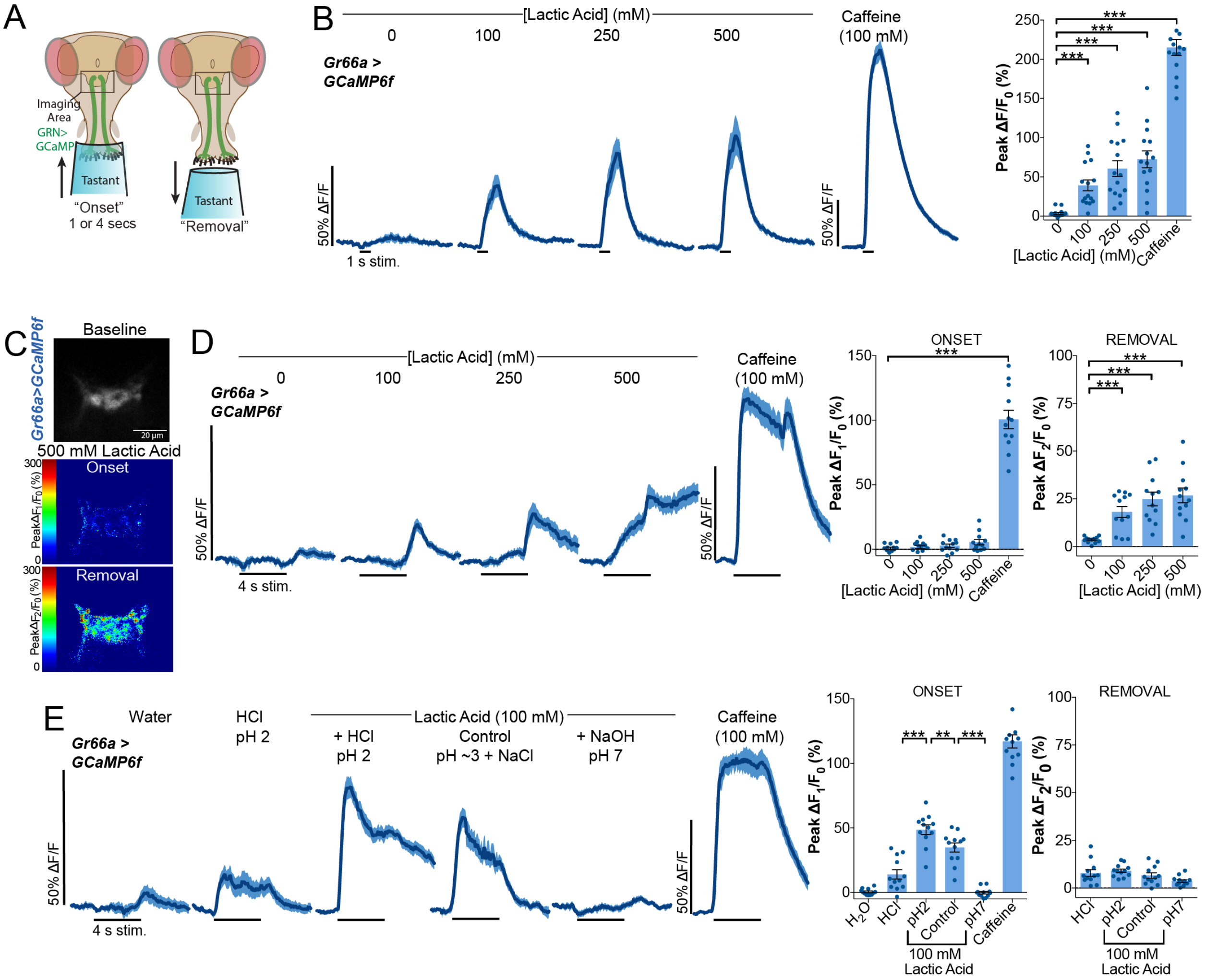


**Figure S3, related to Figure 3:**
**pH influences lactic acid responses in bitter GRNs**
(**A**) Schematic of *in vivo* calcium imaging preparation. Labellar taste neurons are stimulated with tastant while GCaMP6f fluorescence is recorded in the synaptic terminals in the SEZ region of the brain. Tastant is left over the labellum for either 1 or 4 s. (**B**) Bitter GRN calcium responses to 1-second stimulation with lactic acid, in flies expressing GCaMP6f under control of *Gr66a-Gal4*. Lines and shaded areas represent mean ±SEM over time (left). Peak fluorescence changes during each stimulation (right). Bars represent mean ±SEM. n=15 flies. Asterisks denote significant difference from negative control (0 mM, water) by one-way ANOVA with Dunnett's post test, ***p<.001. (**C**) Representative heat map showing *Gr66a>GCaMP6f* fluorescence changes with 500 mM lactic acid stimulus onset vs. removal. (**D**) Bitter GRN calcium responses with 4-second stimulations. Lines and shaded areas represent mean ±SEM over time (left). Quantification of 'onset' and 'removal' peak fluorescence changes during each stimulation and removal of stimulus (right). Bars represent mean ±SEM. n=12 flies, each receiving all tastants shown in order. Asterisks denote significant difference from negative control (0 mM, water) by one-way ANOVA with Dunnett's post test, ***p<.001. (**E**) Bitter GRN responses to pH-adjusted 100 mM lactic acid and control solutions. Lines and shaded areas represent mean ±SEM (left). Quantification of 'onset' and 'removal' peak fluorescence changes during each stimulation and removal of stimulus (right). Bars represent mean ±SEM. n=12 flies, each receiving all tastants in random order (except water always first and caffeine last). Asterisks denote significant differences between control 100 mM lactic acid and other test solutions by one-way ANOVA with Sidak's post test, **p<.01,***p<.001.

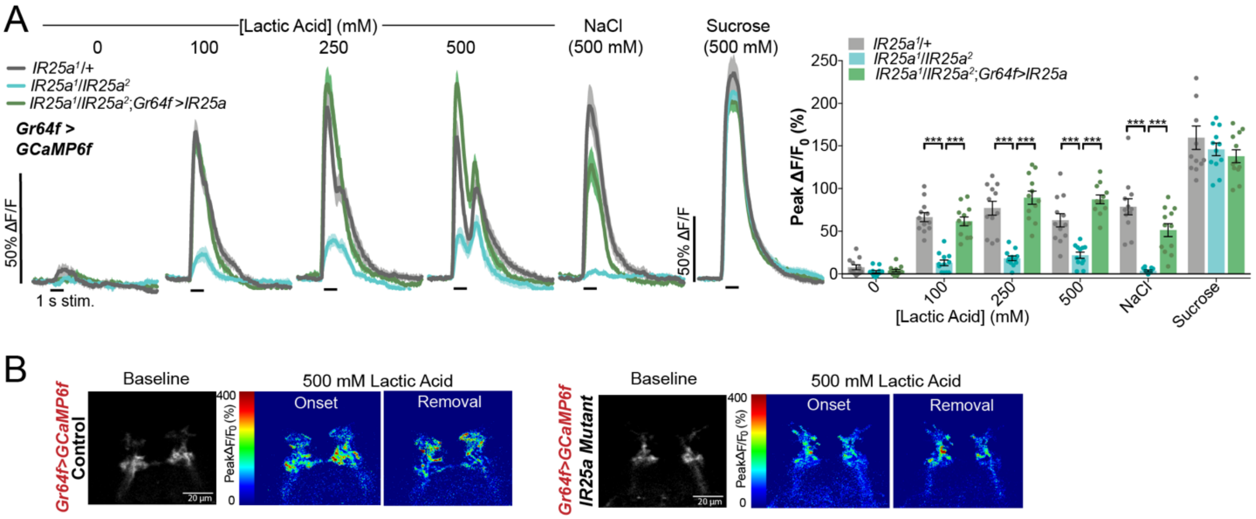

**Figure S4, related to Figure 4:**
**IR25a rescue in sweet GRNs restores calcium imaging responses to lactic acid**
(**A**) Sweet GRN calcium responses in *IR25a* mutants, heterozygous controls, and rescue of IR25a using *Gr64f-Gal4*. Lines and shaded areas represent mean ±SEM over time (left). Quantification of peak fluorescence changes during each (right). Bars represent mean ±SEM. n=12 flies per genotype. Asterisks denote significant differences by two-way ANOVA with Tukey's post test, *p<.05, ***p<.001. (**B**) Representative heat map showing *Gr64f>GCaMP6f* fluorescence changes with 500 mM lactic acid stimulus onset vs. removal in *IR25a* mutant and heterozygous control.

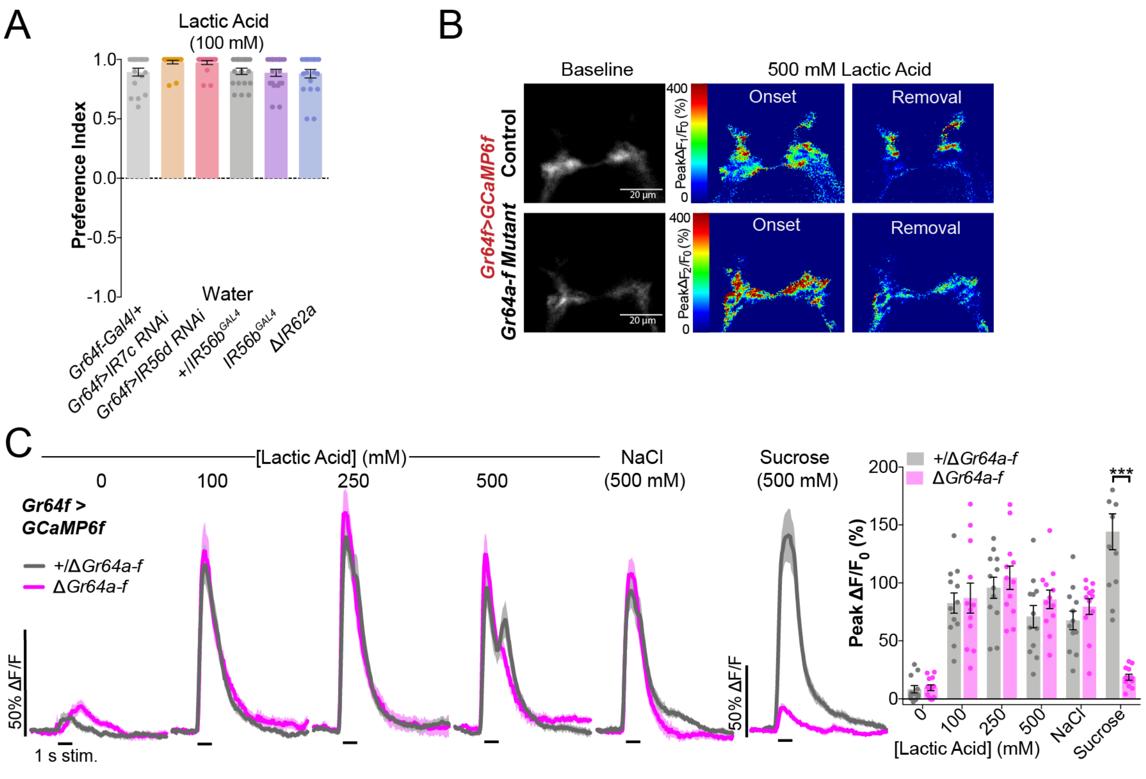


**Figure S5, related to Figure 5:**
**IR behavioural screen and additional imaging in sweet GR mutants**
(**A**) Feeding preference of flies with candidate IRs mutated or knocked down by RNAi. Positive values indicate preference for 100 mM lactic acid; negative values indicate preference for water. Bars represent mean ±SEM. n=18-20 groups of 10 flies per genotype. No significant differences by one-way ANOVA with Tukey's post test. (**B**) Representative heat map showing sweet GRN fluorescence changes with 500 mM lactic acid stimulus onset vs. removal in Δ*Gr64a-f* mutant and heterozygous control. (**C**) Sweet GRN calcium responses in Δ*Gr64a-f* mutants and heterozygous controls with 1-second stimulations. Lines and shaded areas represent mean ±SEM over time (left). Quantification of peak fluorescence changes during each (right). Bars represent mean ±SEM. n=12 flies per genotype. Asterisks denote significant differences by two-way ANOVA with Sidak's post test, ***p<.001.

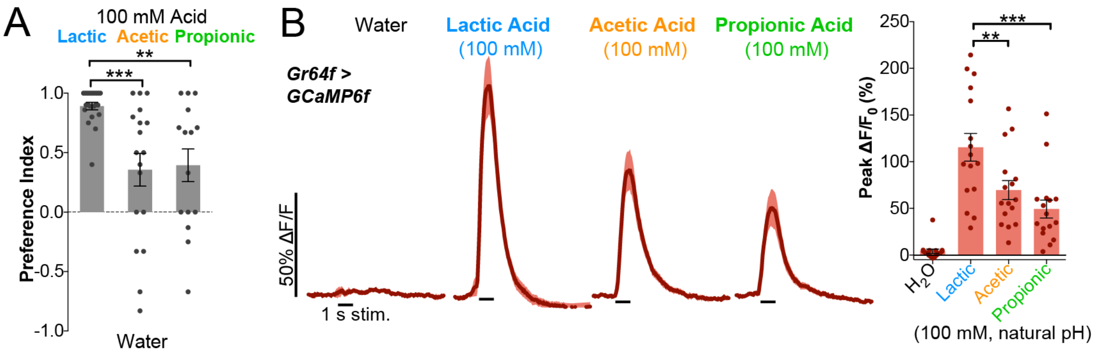


**Figure S6, related to Figure 7:**
**Relative feeding preference and sweet GRN activation with other attractive carboxylic acids**
(**A**) Binary choice feeding preference of $w^{1118}$ flies. Positive values indicate preference for 100 mM indicated acid (lactic, acetic, propionic); negative values indicate preference for water. Bars represent mean ±SEM. n=15-21 groups of 10 flies per group. Asterisks indicate significant differences by one-way ANOVA with Dunnett's post test, **p<.01, ***p<.001. (**B**) Sweet GRN calcium responses in control flies with 1-second stimulations. Lines and shaded areas represent mean ±SEM over time (left). Quantification of peak fluorescence change during each stimulation (right). Bars represent mean ±SEM. n=16 flies. Asterisks indicate significant differences by one-way ANOVA with Dunnett's post test, **p<.01, ***p<.001.
